# Genetic overlap between psychotic experiences in the community across age and with psychiatric disorders

**DOI:** 10.1101/718015

**Authors:** Wikus Barkhuizen, Oliver Pain, Frank Dudbridge, Angelica Ronald

## Abstract

**Background:** This study explores the degree to which genetic influences on psychotic experiences are stable across adolescence and adulthood, and their overlap with psychiatric disorders.

**Methods:** Genome-wide association results were obtained for adolescent psychotic experiences and negative symptom traits (N = 6,297-10,098), schizotypy (N = 3,967-4,057) and positive psychotic experiences in adulthood (N = 116,787-117,794), schizophrenia (N = 150,064), bipolar disorder (N = 41,653) and depression (N = 173,005). Linkage disequilibrium score regression was used to estimate genetic correlations. Implicated genes from functional and gene-based analyses were compared. Mendelian Randomization was performed on trait pairs with significant genetic correlations.

**Results:** Subclinical auditory and visual hallucinations and believing in conspiracies during adulthood were significantly genetically correlated with schizophrenia (r_g_ = .27-.67) and major depression (r_g_ = .41-.96) after correction for multiple testing. Auditory and visual subclinical hallucinations were highly genetically correlated (r_g_ = .95). Cross-age genetic correlations for psychotic experiences were not significant. Gene mapping and gene association analyses revealed 14 possible genes associated with psychotic experiences that overlapped across age for psychotic experiences or between psychotic experiences and psychiatric disorders. Mendelian Randomization indicated bidirectional associations between auditory and visual hallucinations in adults but did not support causal relationships between psychotic experiences and psychiatric disorders.

**Conclusions:** Psychotic experiences in adulthood may be more linked genetically to schizophrenia and major depression than psychotic experiences in adolescence. Our study implicated specific genes that are associated with psychotic experiences across development as well as genes shared between psychotic experiences and psychiatric disorders.

## Introduction

Psychotic experiences (also called *‘psychotic-like experiences’)* resemble positive symptoms of psychotic disorders, like paranoia, hallucinations and cognitive symptoms, and can be measured in the general population. Negative symptom traits in the community resemble negative symptoms of psychotic disorders such as apathy, anhedonia and social withdrawal. Schizotypy (1) is a related, older personality-based construct compared to psychotic experiences and negative symptoms (PENS). Positive psychotic experiences during adolescence or adulthood, especially when persistent, are associated with an increased risk of developing psychotic disorders (2-7) and to a lesser extent other psychiatric disorders (8-10). Likewise, schizotypy is associated with subsequently developing psychotic disorders (11).

Twin studies suggest heritability accounts for a third to a half of variation in PENS during adolescence (12-17) and recent studies suggest modest single nucleotide polymorphism heritability (SNP-h^2^) for some PENS in mid-adolescence (3-9%) and for schizotypy in adults (20-27%) (18-20). Evidence from genome-wide association studies (GWAS) indicate that PENS during adolescence share genetic influences with schizophrenia and negative symptom traits with major depression (18, 21-23), although other studies did not find this, particularly those that used comparatively smaller samples or polygenic scores (PGS) from less well-powered GWAS (21, 23-26). A study on adults reported an association between schizophrenia PGS and schizotypy assessed using semi-structured interviews, but not with self-rated PENS (27). However other adult samples show no association between genome-wide significant schizophrenia variants and social anhedonia (28) and no positive associations between schizophrenia PGS and adult psychotic experiences (29). These studies calculated PGS from GWAS based on smaller samples than currently available. In sum, newer studies using more reliable PGS tend to report significant genetic overlap between PENS and psychiatric disorders. However, studies have mainly focussed on adolescents and young adults rather than older adults and have not reported on the genetic overlap of psychotic experiences across age, despite this being an important topic for understanding the etiology and development of mental illness. It is also not known if psychotic experiences and schizotypy share genetic influences.

This study investigates the genetic overlap between psychotic experiences across age using the largest current GWAS summary statistics available for adolescent PENS, and for schizotypy and positive psychotic experiences measured in adulthood. We evaluate genome-wide genetic correlations and overlapping associated genes between these trait measures and with schizophrenia, depression and bipolar disorder. For traits and disorders that share common additive genetic influences, we further explore the nature of these associations using Mendelian Randomization.

## Methods and Materials

### Samples and measures

#### Psychotic experiences and negative symptom traits during adolescence

Summary statistics for adolescent PENS came from a mega-GWAS of three European community samples consisting of 6,297-10,098 individuals (18): TEDS (Twins Early Developmental Study)(30), a community sample born between 1994-1996 in England and Wales (mean age 16.32 years); ALSPAC (Avon Longitudinal Study of Parents and Children)(31, 32), a birth cohort from the United Kingdom (mean age 16.76 years) born in 1991-1992; and CATSS (Child and Adolescent Twin Study in Sweden)(33) that recruited twins born in Sweden since 1992 (mean age 18.31 years). Additional information is provided in the supplementary note.

In TEDS, PENS items came from the Specific Psychotic Experiences Questionnaire (34) and were matched by a team of clinicians to similar items from psychopathology questionnaires available in CATSS and ALSPAC (18). After harmonisation, PENS included four continuous subscales that assessed the frequency or severity of paranoia and hallucinations, cognitive disorganization, anhedonia and parent-rated negative symptoms.

#### Schizotypy during adulthood

Schizotypy was assessed in the Northern Finland Birth Cohort 1996 (NFBC)(35) when participants were aged 31 years. GWAS summary statistics (20) of four continuous scales of schizotypy were included (N = 3,967 – 4.057): Perceptual aberrations were assessed with the Perceptual Aberration Scale (36) and included experiences that resemble clinical features of schizophrenia. Hypomania was from the Hypomanic Personality Scale (37), devised to assess hypomania, gregariousness, grandiosity and euphoria. Two scales from Chapman’s Schizotypia Scales were employed: The Revised Social Anhedonia Scale and the Revised Physical Anhedonia Scale (38), devised to assess the inability to take pleasure from physical and social stimuli, respectively.

#### Positive psychotic experiences assessed in adults

GWAS summary statistics of four dichotomous items from the UK Biobank were obtained from Neale Lab (*http://www.nealelab.is/uk-biobank*) for individuals of European ancestry (N = 116,787 - 117,794). Items assessed psychotic experiences in adults aged 40-69 years: Whether participants ever experienced hearing an un-real voice, had an un-real vision, held a belief in an un-real conspiracy and experienced un-real communications or signs.

Additional information on psychotic experiences items are provided in the Supplementary Note.

#### Psychiatric disorders

GWAS summary statistics for schizophrenia (39), bipolar disorder (40) and major depressive disorder (41) were downloaded from the Psychiatric Genetics Consortium (https://www.med.unc.edu/pgc/results-and-downloads). Summary statistics for depression excluded 23andMe participants.

### Analyses

Quality control procedures were applied to summary statistics prior to analyses. Genetic variants were removed if they had incomplete association statistics, were non-biallelic, or strand ambiguous. Variants with alleles that did not match those in the 1000 Genomes reference panel (phase 3), with info scores < .9 and minor allele frequency (MAF) < .01 were excluded.

Linkage disequilibrium (LD) score regression (42, 43) was used to estimate SNP-h^2^ for traits and the genetic correlation (r_g_) between traits. Variants were merged with the HapMap3 reference panel as recommended to ensure good imputation quality (42). Heritability and genetic correlations were converted to liability scales using lifetime prevalence of 1% for schizophrenia, 15% for depression and 2% for bipolar disorder (44-46). Effective sample sizes were used for adolescent psychotic experiences because TEDS and CATSS included siblings (for details, see (18)). For genetic correlations where there was no known sample overlap, the LD score regression intercept was constrained to zero, but left unconstrained between positive psychotic experiences and depression that both included UK Biobank participants, and between psychiatric disorders that include PGC participants. Benjamini-Hochberg correction for multiple testing was performed at a false discovery rate (FDR) of 0.05 to account for 105 correlations estimated.

Gene-wide associations and gene mapping were performed using the FUMA pipeline (46) and results were compared across psychotic experiences and psychiatric disorders. To identify genes at genome-wide significance, MAGMA vl.06 (48) was used to aggregate the *p-* values of SNPs within gene coding regions (0kb annotation window). Default parameters and a SNP-wide mean model was employed. Bonferroni corrected p-value thresholds were set for the number of genes tested within each phenotype. Positional mapping of variants within 10kb of gene regions and likely to have functional consequences (CADD score ≥ 12.37) and gene mapping using expression quantitative trait loci (eQTL) associations and chromatin interactions was performed for independent lead SNPs (at p < 1 × 10^-5^ for psychotic experiences, p < 1 × 10^-6^ for depression, and p < 1 × 10^-8^ for schizophrenia and bipolar disorder) and 1000 Genomes reference panel variants in LD with independent SNPs at r2 ≥ 0.6 using the recommended parameters in FUMA(47). Additional details are provided in the Supplementary Note.

Mendelian randomization (MR)(49) was conducted to test for causal relationships between phenotypes that had significant genetic correlations. SNPs were selected as instrumental variables based on the clumping algorithm in PLINK (50) using an r^2^ threshold of .05 within a 500kb window size at genome-wide significant levels (p < 5 × 10^-8^) for schizophrenia. Summary statistics for depression did not include 23andMe participants and had an insufficient number of genome-wide significant variants for MR analyses. Instead, genome-wide significant variants were obtained from the publication (41). Insufficient genome-wide significant variants associated with psychotic experiences meant that p-value thresholds for psychotic experiences were set at *p* < 5 × 10^-5^ to allow for at least 20 independent SNPs in MR analysis.

Generalised Summary-data-based Mendelian Randomization (GSMR) was used due to its advantages of accounting for residual LD structure between instrumental variables (set at LD r^2^ > 0.1) and accounting for sampling variation in the exposure and outcome GWAS (51). MR-Egger regression (52), Weighted Median (53) and Weighted Mode (54) methods were conducted as sensitivity analyses for possible violation of MR assumptions. Potentially pleiotropic SNPs identified by Heidi-outlier analysis and SNPs excluded due to residual LD structure during GSMR analysis were also excluded from sensitivity analyses.

## Results

Common additive genetic variance accounted for 8-10% of phenotypic variation in adolescent PENS (paranoia/hallucinations and negative symptoms did not have significant SNP-h^2^ estimates in these analyses), 30-37% in schizotypy during adulthood and 7-10% in positive PE during adulthood (Table 1).

**Table 1.** Genome wide association study sample sizes and SNP-heritability estimates.

| | GWAS N | N cases | QC-positive SNPs | SNP-$h^2$ | SE | *p* |
| --- | --- | --- | --- | --- | --- | --- |
| **Adolescent psychotic experiences and negative symptom traits (18)** |  |  |  |  |  |  |
| Paranoia and hallucinations | 8,665 | *Continuous* | 3,363,829 | -0.0042 | 0.0352 | .453 |
| Cognitive disorganization | 6,297 | *Continuous* | 3,363,829 | 0.1048 | 0.0566 | .032 |
| Anhedonia | 6,579 | *Continuous* | 3,363,829 | 0.0797 | 0.0479 | .048 |
| Parent-rated negative symptoms | 10,098 | *Continuous* | 3,363,829 | -0.0222 | 0.0316 | .241 |
| **Schizotypy during adulthood (34)** |  |  |  |  |  |  |
| Hypomania | 3,967 | *Continuous* | 5,493,986 | 0.3732 | 0.1011 | <.001 |
| Perceptual aberrations | 4,057 | *Continuous* | 5,493,986 | 0.3037 | 0.0916 | <.001 |
| Physical anhedonia | 3,988 | *Continuous* | 5,493,986 | 0.3655 | 0.0965 | <.001 |
| Social anhedonia | 4,025 | *Continuous* | 5,493,986 | 0.2950 | 0.0826 | <.001 |
| **Positive psychotic experiences (Neale Lab, 2018)** |  |  |  |  |  |  |
| Un-real voice | 117,503 | 2,009 | 6,443,634 | 0.0709 | 0.0255 | .003 |
| Un-real vision | 116,787 | 3,768 | 6,443,706 | 0.1032 | 0.0224 | <.001 |
| Un-real conspiracy | 117,794 | 932 | 6,443,695 | 0.0910 | 0.0521 | .040 |
| Un-real communications or signs | 117,731 | 822 | 6,443,693 | 0.0666 | 0.0499 | .091 |
| **Psychiatric disorders** |  |  |  |  |  |  |
| Schizophrenia (38) | 150,064 | 36,989 | 5,274,747 | 0.1631 | 0.0044 | <.001 |
| Bipolar disorder (39) | 41,653 | 20,129 | 5,083,505 | 0.2999 | 0.0102 | <.001 |
| Major Depression (40) | 173,005 | 59,851 | 5,488,968 | 0.0999 | 0.0042 | <.001 |

*Note:* SNP- $h^2$ = univariate SNP heritability; SNP- $h^2$ converted to a liability scale for binary traits; Effective sample size used for adolescent PENS in LD score regression analyses to account for the presence of siblings (paranoia and hallucinations = 7970.416; cognitive disorganization = 5082.760; anhedonia = 6068.311; parent-rated negative symptoms = 8763.295).

Genetic correlations are shown in Figure 1. Note that in pairwise comparisons with paranoia and hallucinations and parent-rated negative symptoms, genetic correlations could not be computed, likely due to low SNP-h^2^. For these comparisons, we report genetic covariance (*ρ*_g_) which indicates direction of correlation (Table S1).

**Figure 1.**
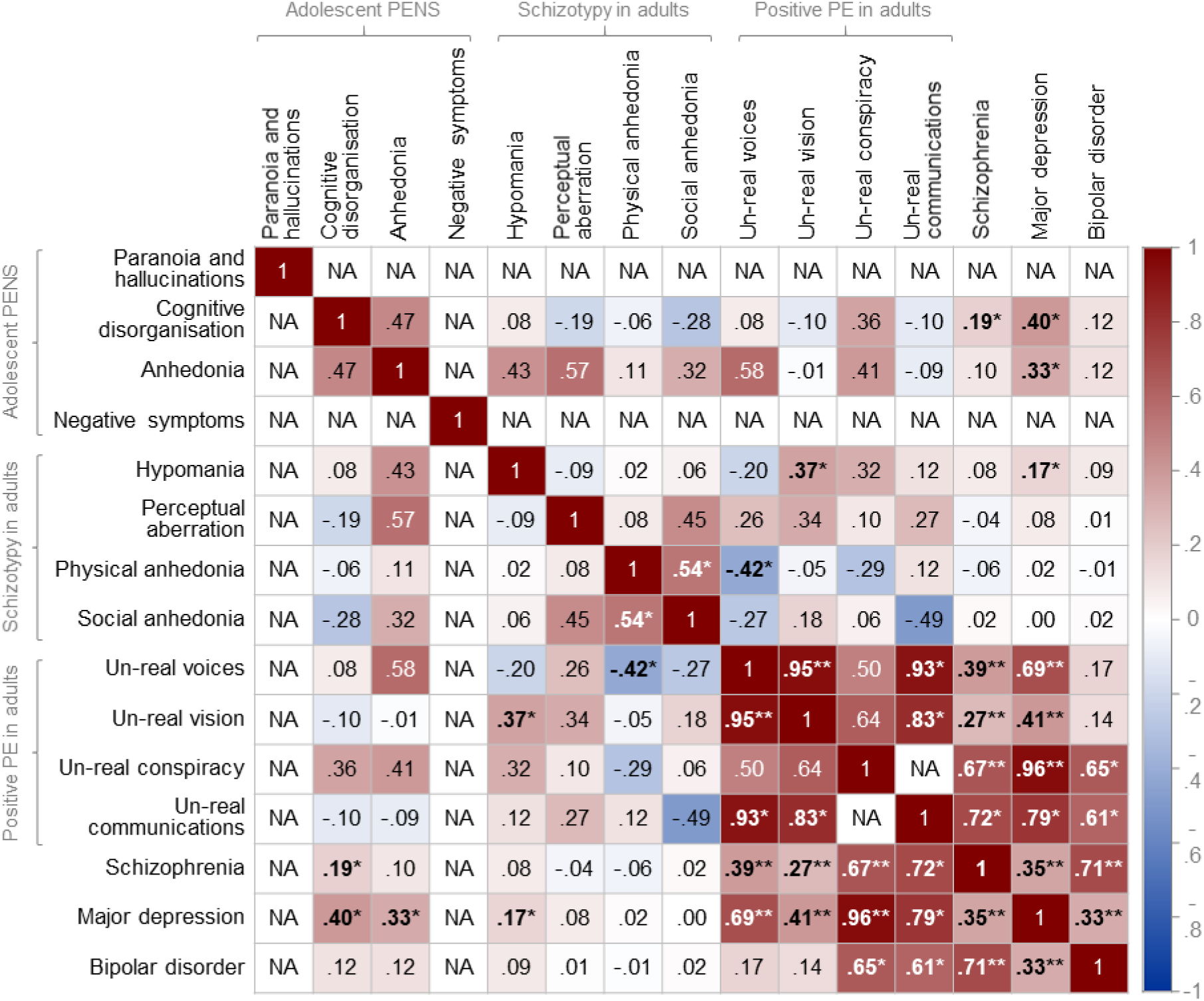
Heat map showing genetic correlations between psychotic experiences and psychiatric disorders. *Note:* PENS = Psychotic experiences (PE) and negative symptom traits; NA = genetic correlations could not be computed due to low SNP-heritability or sample size (see Table S1 for genetic covariance estimates); ⋆ indicates nominally statistically significant genetic correlations at p<.05; ⋆⋆ indicates genetic correlations that survived Benjamini-Hochberg correction for multiple testing for 105 pairwise correlations (at a FDR of 0.05); Genetic correlations reported using unconstrained LD score regression intercept between phenotypes with sample overlap.

### Genetic correlations with psychiatric disorders

For positive psychotic experiences in the UK Biobank, hearing un-real voices was significantly genetically correlated with schizophrenia (r_g_= .39, p = 2.27 × 10^-5^ and depression (r_g_= .69, p = 8.07 × 10^-6^ but not with bipolar disorder (r_g_= .17, p = .082). Likewise, un-real visions was significantly correlated with schizophrenia (r_g_= .27, p = 4.12 × 10^-7^ and depression (r_g_ = .41, p = 2.00 × 10^-4^ but not with bipolar disorder (r_g_= .14, p = .149). We observed high and significant genetic correlations between un-real conspiracies with schizophrenia (r_g_= .67, *p* = .002) and depression (r_g_= .96, p = .001) while the genetic correlation with bipolar disorder was not significant after correction for multiple testing (r_g_= .65, p = .019). Genetic correlations between un-real communications with schizophrenia (r_g_= .72, p = .007), depression (r_g_= .79, p = .021) and bipolar disorder (r_g_= .61, p = .033) were at nominal significance (p < .05) but did not survive correction for multiple testing.

Between PENS during adolescence and psychiatric disorders, genetic correlations were not significant after correction for multiple testing. Nominally significant genetic correlations were observed between cognitive disorganization with schizophrenia (r_g_ = .19, p = .034) and depression (r_g_ = .40, p = .006) and between anhedonia and depression (r_g_ = .33, p = .021). Negative symptoms covaried positively with schizophrenia (*ρ*_g_ *=* .03, p = 1.37 × 10^-4^ and depression (*ρ*_g_ *=* .03, p = 2.41 × 10^-7^ Positive genetic covariation was observed between paranoia and hallucinations and depression (*ρ*_g_ *=* .03, p = 1.43 × 10^-4^ No significant genetic correlations were found between adolescent PENS and bipolar disorder.

Between psychiatric disorders and schizotypy in adults, hypomania was genetically correlated with depression at nominal significance (r_g_ = .21, p = .016). We did not find evidence that the schizotypy scales correlated with schizophrenia or bipolar disorder.

### Genetic stability of psychotic experiences across age

PENS during adolescence were not significantly genetically correlated with positive psychotic experiences and schizotypy during adulthood. Between the adult samples, hypomania was genetically correlated with un-real visions (r_g_ = .37, p = .024) and physical anhedonia showed a negative genetic correlation with un-real voices (r_g_ = -.42, p = .034), but not after correction for multiple testing.

### Genetic correlations within samples

Between positive psychotic experiences items within the UK Biobank, un-real voices and un-real visions were highly and significantly genetically correlated (r_g_= .95, p = 7.61 × 10^-5^ Genetic correlations between un-real communications with un-real voices (r_g_= .93, p = .038) and visions (r_g_= .83, p = .050) were not significant after correction for multiple testing. Un-real conspiracies were not genetically correlated with un-real voices or visions. A genetic correlation between un-real conspiracies and un-real communications could not be computed due to low SNP-h^2^ in both phenotypes but these two items showed positive genetic covariance (*ρ*_g_ *=* .01, p = .003).

There was a nominally significant association between physical and social anhedonia (r_g_= .54, p = .044). All other genetic correlations between the schizotypy scales in the North Finland Birth Cohort were not significant.

Between the adolescent PENS scales, a positive genetic covariance was identified between anhedonia and parent-rated negative symptoms (*ρ*_g_ *=* .11, p = .012). No significant genetic overlap between other pairs of adolescent PENS was found.

### Comparison of implicated genes across mapping strategies and phenotypes

Gene mapping and genome-wide gene association results from FUMA were compared across psychotic experiences and psychiatric disorders (Table 2). Full results for each phenotype are provided in Tables S2-7 and Figures Sl-3. Results revealed 32 genes for adolescent PENS, of which *PAN3* mapped to cognitive disorganisation in adolescence and schizotypy in adulthood (perceptual aberrations) and *NADK2* to negative symptoms in adolescence and positive psychotic experiences in the UK Biobank (un-real communications); None overlapped with psychiatric disorders. Seventy-three genes were found for adult schizotypy including the aforementioned *PAN3* and six genes that were also indicated for schizophrenia. *ANK3* overlapped with both positive psychotic experiences (un-real visions) and schizophrenia. For positive psychotic experiences in adults, 104 genes were identified of which seven genes overlapped with schizophrenia and as mentioned above, one with schizotypy and one with adolescent PENS (Figure 2).

**Table 2.** Gene-associations and gene-mapping results for overlapping genes across phenotypes.

| Gene | Chr | Phenotype | MAGMA | | Positional mapping | | eQTL mapping | | | Chromatin Interactions | Mapped SNPs<br>min $p_{\text{GWAS}}$ | Independent lead SNPs |
| --- | --- | --- | --- | --- | --- | --- | --- | --- | --- | --- | --- | --- |
| | | | $n$ SNPs | $p_{\text{MAGMA}}$ | $n$ SNPs | CADD | $n$ SNPs | GTEx map | Min $p_{\text{eQTL}}$ | HiCi cortex | | |
| ANK3 | 10 | Physical anhedonia | - | - | 4 | 15.60 | - | - | - | - | $6.35 \times 10^{-6}$ | rs72818480 |
| | | Unreal visions | - | - | 1 | 13.71 | - | - | - | - | $9.71 \times 10^{-6}$ | rs10994279 |
| | | Schizophrenia | 1580 | $9.46 \times 10^{-7}$ | - | - | - | - | - | - | - | - |
| CACNA1C | 12 | Hypomania | - | - | 3 | 17.74 | 85 | CerHem;<br>Cereb | $7.34 \times 10^{-9}$ | - | $1.91 \times 10^{-6}$ | rs6489351; rs3922316;<br>rs34382810 |
| | | Schizophrenia | 1295 | $7.79 \times 10^{-20}$ | 4 | 17.74 | 86 | CerHem;<br>Cereb | $7.34 \times 10^{-9}$ | - | $2.63 \times 10^{-17}$ | rs6489351; rs11062159;<br>rs12423277; rs2239037;<br>rs2007044; rs882193; rs4765914;<br>rs2239063 |
| CLK1 | 2 | Perceptual aberrations | - | - | - | - | - | - | - | Fetal | $4.65 \times 10^{-6}$ | rs56225831 |
| | | Schizophrenia | - | - | - | - | - | - | - | Adult | $3.60 \times 10^{-13}$ | rs796364; rs74938253 |
| DCP1B | 12 | Hypomania | - | - | - | - | - | - | - | Fetal/adult | $2.16 \times 10^{-6}$ | rs34382810; rs3922316;<br>rs6489351 |
| | | Schizophrenia | - | - | - | - | - | - | - | Fetal/adult | $1.65 \times 10^{-16}$ | rs2007044; rs11062159;<br>rs2239037; rs12423277;<br>rs6489351; rs2239063 |
| FAM216A | 12 | Unreal visions | - | - | - | - | - | - | - | Fetal | $1.55 \times 10^{-6}$ | rs4766567 |
| | | Schizophrenia | - | - | - | - | - | - | - | Fetal | $7.09 \times 10^{-10}$ | rs4766428 |
| FKBP4 | 12 | Hypomania | - | - | - | - | - | - | - | Fetal/adult | $2.16 \times 10^{-6}$ | rs34382810; rs3922316 |
| | | Schizophrenia | - | - | - | - | - | - | - | Fetal/adult | $1.65 \times 10^{-16}$ | rs2007044; rs12423277;<br>rs11062159; rs2239037 |
| GPN3 | 12 | Unreal visions | - | - | - | - | - | - | - | Fetal | $1.55 \times 10^{-6}$ | rs4766567 |
| | | Schizophrenia | - | - | - | - | - | - | - | Fetal | $7.09 \times 10^{-10}$ | rs4766428 |
| NADK2 | 5 | Negative symptoms | - | - | - | - | - | - | - | Fetal/adult | NA<sup>a</sup> | rs147205145 |
| | | Unreal communications | - | - | - | - | - | - | - | Adult | $3.68 \times 10^{-6}$ | rs16902775 |
| PAN3 | 13 | Cognitive disorganization | - | - | 1 | 14.24 | - | - | - | - | NA<sup>a</sup> | rs9513058 |
| | | Perceptual aberrations | - | - | - | - | - | - | - | Adult | $4.92 \times 10^{-3}$ | rs9581857 |
| PPP1CC | 12 | Unreal visions | - | - | - | - | - | - | - | Fetal | $1.55 \times 10^{-6}$ | rs4766567 |
| | | Schizophrenia | - | - | - | - | - | - | - | Fetal | $7.09 \times 10^{-10}$ | rs4766428 |

**Table 2.** (continued).
| Gene | Chr | Phenotype | MAGMA | | Positional mapping | | eQTL mapping | | | Chromatin Interactions | Mapped SNPs<br>min $p_{\text{GWAS}}$ | Independent lead SNPs |
| --- | --- | --- | --- | --- | --- | --- | --- | --- | --- | --- | --- | --- |
| | | | $n$ SNPs | $p_{\text{MAGMA}}$ | $n$ SNPs | CADD | $n$ SNPs | GTEx map | Min $p_{\text{eQTL}}$ | HiCi cortex | | |
| SNX7 | 1 | Unreal visions | - | - | - | - | - | - | - | Fetal | $3.78 \times 10^{-4}$ | rs34186519 |
| | | Unreal conspiracy | - | - | - | - | - | - | - | Fetal | $4.42 \times 10^{-8}$ | rs4908317; rs17403033 |
| | | Schizophrenia | - | - | - | - | - | - | - | Fetal/adult | $2.83 \times 10^{-17}$ | rs1702294; rs2802525;<br>rs61786697; rs7526108 |
| TMEM116 | 12 | Unreal visions | - | - | - | - | 2 | BasalG | $7.86 \times 10^{-7}$ | - | $1.76 \times 10^{-5}$ | rs4766567 |
| | | Schizophrenia | - | - | - | - | - | - | - | Fetal | $7.09 \times 10^{-10}$ | rs4766428 |
| TSNARE1 | 8 | Unreal communications | - | - | - | - | 3 | CerHem | $7.77 \times 10^{-7}$ | Fetal/adult | $5.64 \times 10^{-5}$ | rs10102944 |
| | | Schizophrenia | 502 | $5.26 \times 10^{-10}$ | 1 | 13.06 | - | - | - | Fetal/adult | $2.85 \times 10^{-13}$ | rs4129585; rs74576444;<br>rs62512616; rs13282237;<br>rs72687362 |
| TSPAN9 | 12 | Hypomania | - | - | - | - | - | - | - | Fetal/adult | $2.16 \times 10^{-6}$ | rs34382810; rs3922316 |
| | | Schizophrenia | - | - | - | - | - | - | - | Fetal/adult | $1.65 \times 10^{-16}$ | rs2007044; rs12423277;<br>rs11062159; rs2239037;<br>rs2239063 |

*Note:* Chr = chromosome number; CADD = The maximum Combined Annotation-Dependent Depletion score; Min $p_{\text{GWAS}}$ = minimum p-value of annotated or mapped variants from summary statistics; CerHem = GTEx v7 Cerebellar hemisphere tissue map; Cereb = GTEx v7 cerebellum tissue map; BasalG = GTEx v6 Basal ganglia tissue map. SNPs mapped to genes included independent lead SNPs and SNPs from 1K Genomes reference panel in LD with lead SNPs ( $r^2 \geq 0.6$ ), within 10kb of locus and likely to be deleterious (CADD $\geq 12.37$ ).
<sup>a</sup> Annotated or mapped variants present only in reference panel and therefore does not have a minimum $p_{\text{GWAS}}$ .

**Figure 2.**
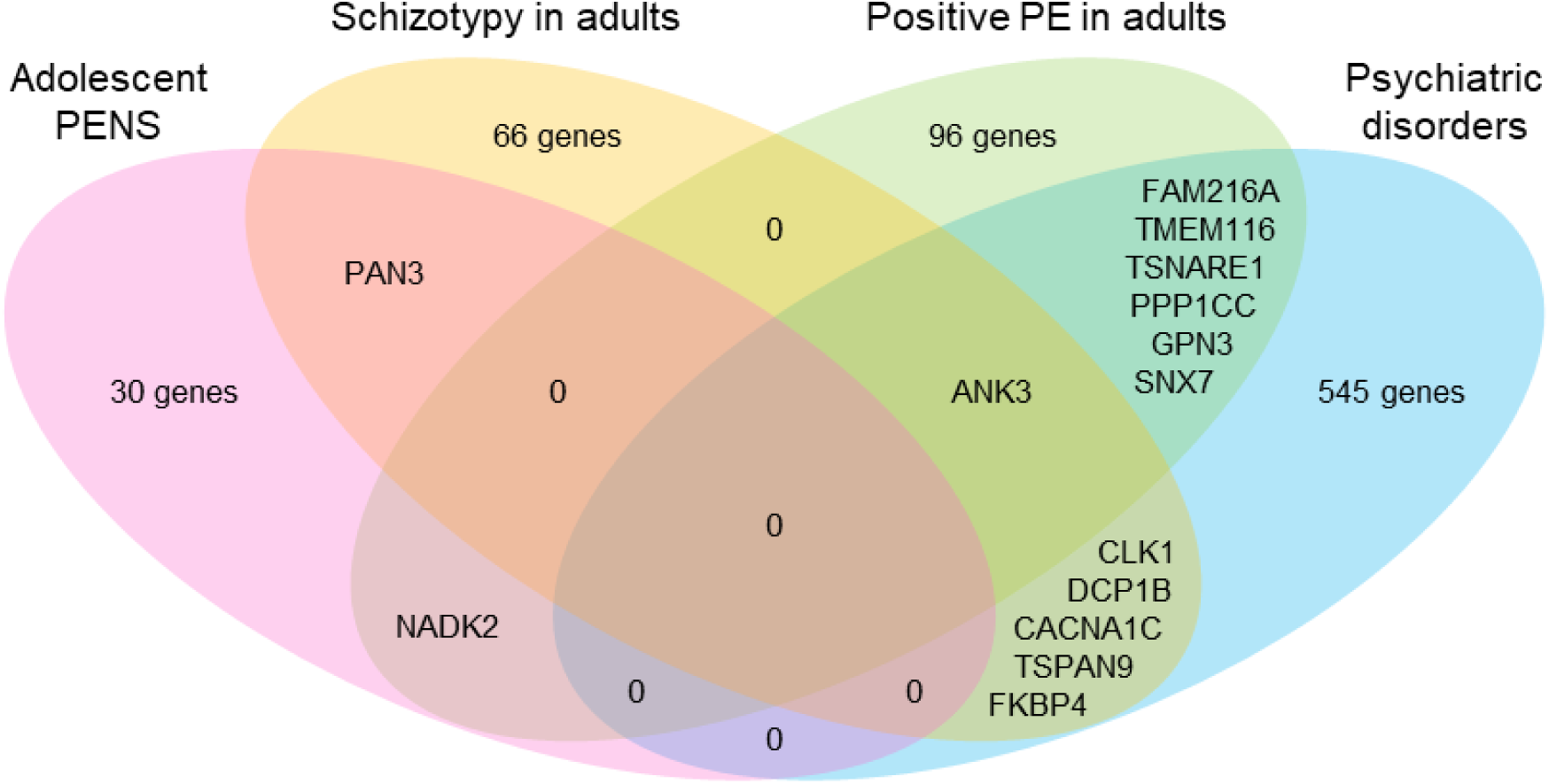
Number of overlapping genes between psychotic experiences across age and psychiatric disorders. *Note:* PENS = Psychotic experiences and negative symptom traits. Genes identified using a) genome-wide gene associations in MAGMA after Bonferroni correction for the number of gene associations tested, b) positional mapping that prioritized genes based on variant functional annotations obtained using ANNOVAR, c) eQTL (expressive quantitative trait) mapping and d) chromatin interaction mapping.

### Mendelian randomization

Results from GSMR analyses and MR sensitivity analyses are summarised in Table 3 and MR-Egger intercept tests, heterogeneity statistics and plots in Table S8 and Figures S4-5.

**Table 3.** Mendelian randomization analyses.

| Exposure | Outcome | Heidi<br>SNPs | LD<br>SNPs | n<br>SNP | GSMR results |  |  | MR-Egger |  |  | Weighted Median |  |  | Weighted Mode |  |  |
| --- | --- | --- | --- | --- | --- | --- | --- | --- | --- | --- | --- | --- | --- | --- | --- | --- |
|  |  |  |  |  | Beta | SE | *p* | Beta | SE | *p* | Beta | SE | *p* | Beta | SE | *p* |
| Un-real voice | → Schizophrenia | 0 | 0 | 49 | 0.021 | 0.676 | 0.975 | 2.494 | 1.996 | 0.218 | -0.008 | 0.908 | 0.993 | -0.216 | 1.993 | 0.914 |
| Schizophrenia | → Un-real voice | 0 | 11 | 98 | 0.001 | 0.001 | 0.281 | 0.001 | 0.004 | 0.846 | 0.001 | 0.001 | 0.424 | 0.003 | 0.003 | 0.281 |
| Un-real visions | → Schizophrenia | 0 | 0 | 48 | -0.115 | 0.501 | 0.819 | -1.860 | 1.697 | 0.279 | -0.132 | 0.707 | 0.852 | -1.091 | 1.635 | 0.508 |
| Schizophrenia | → Un-real visions | 2 | 11 | 96 | 0.002 | 0.001 | 0.084 | -0.003 | 0.005 | 0.514 | 0.003 | 0.002 | 0.099 | 0.005 | 0.004 | 0.270 |
| Un-real voice | → Major depression | 0 | 0 | 55 | -0.269 | 0.488 | 0.581 | 1.025 | 1.405 | 0.469 | 0.091 | 0.685 | 0.895 | 0.502 | 1.626 | 0.759 |
| Major depression | → Un-real voice | 0 | 0 | 36 | 0.000 | 0.000 | 0.424 | 0.007 | 0.003 | **0.042** | 0.000 | 0.000 | 0.081 | 0.000 | 0.000 | 0.140 |
| Un-real voice | → Un-real visions | 0 | 0 | 97 | 0.321 | 0.034 | **1.73E-21** | 0.310 | 0.071 | **3.47E-05** | 0.333 | 0.046 | **5.16E-13** | 0.505 | 0.133 | **2.66E-04** |
| Un-real visions | → Un-real voice | 0 | 0 | 85 | 0.166 | 0.019 | **4.25E-18** | 0.123 | 0.043 | **0.005** | 0.168 | 0.026 | **1.38E-10** | 0.211 | 0.065 | **1.62E-03** |
| Un-real visions | → Major depression | 0 | 0 | 52 | 0.263 | 0.362 | 0.468 | -0.711 | 1.036 | 0.496 | 0.337 | 0.524 | 0.520 | 0.376 | 1.253 | 0.765 |
| Major depression | → Un-real visions | 0 | 0 | 36 | 0.000 | 0.000 | 0.986 | 0.008 | 0.005 | 0.103 | 0.000 | 0.000 | 0.834 | 0.000 | 0.000 | 0.576 |
| Un-real conspiracy | → Major depression | 0 | 0 | 49 | 0.356 | 0.742 | 0.631 | -1.264 | 2.031 | 0.537 | 1.211 | 1.088 | 0.266 | 1.780 | 2.381 | 0.458 |
| Major depression | → Un-real conspiracy | 0 | 0 | 36 | 0.000 | 0.000 | 0.857 | 0.007 | 0.002 | **0.003** | 0.000 | 0.000 | 0.768 | 0.000 | 0.000 | 0.814 |
| Un-real conspiracy | → Schizophrenia | 0 | 0 | 46 | -0.020 | 1.025 | 0.984 | 1.976 | 3.466 | 0.572 | -0.401 | 1.482 | 0.787 | -0.219 | 2.835 | 0.939 |
| Schizophrenia | → Un-real conspiracy | 1 | 11 | 97 | 0.002 | 0.001 | **9.33E-05** | 0.001 | 0.003 | 0.810 | 0.002 | 0.001 | **0.006** | 0.002 | 0.002 | 0.397 |

Note: GSMR = Generalized Summary-based Mendelian Randomization; LD SNPs = SNPs with residual LD at $r^2 > 0.1$ removed from analysis; n SNPs = number of variants remaining in analyses after those identified as Heidi-outliers or with residual LD was removed.
SNPs identified as having residual LD and as Heidi outliers were also excluded from MR Egger, Weighted Median and Weighted Mode analyses.
Bold text indicates significant *p*-values at $p < 0.05$ .

GSMR analyses provided evidence for a bi-directional association between un-real voices and un-real visions (P_GSMR_ = 1.73 × 10^-21^ for un-real voices as the exposure; P_GSMR_ = 4.25 × 10^-25^ for un-real visions as the exposure) and replicated in all MR sensitivity analyses.

We observed evidence of a directional effect of schizophrenia liability on believing in un-real conspiracies (P_GSMR_ = 9.33 × 10^-5^) but the effect was small and replicated in Weighted Median but not in MR-Egger nor Weighted Mode analyses. No evidence of a significant effect in the other direction was found.

Only the MR-Egger method indicated significant directional effects of a liability to depression on a propensity to report un-real voices and un-real conspiracies. However, MR-Egger intercept tests indicated the presence of directional pleiotropy in both instances (Table S8), indicating that MR-Egger results may be reliable as it adjusts for non-zero intercepts allowing for more robust estimates when horizontal pleiotropy is present compared to the other methods. The discrepancy between MR methods may also be due to outliers not identified by Heidi-outlier analysis.

## Discussion

This study investigated whether common genetic variation associated with specific types of psychotic experiences and negative symptom traits (PENS) overlaps across adolescence and adulthood and with psychiatric disorders. Our results suggest that with increasing age from adolescence to adulthood, psychotic experiences become more linked genetically to schizophrenia and major depression. We did not observe evidence of significant genetic stability of psychotic experiences across adolescence and adulthood in existing data. However, our study implicated specific genes that are associated with psychotic experiences across development as well as genes shared between psychotic experiences and psychiatric disorders.

### Associations between psychotic experiences and psychiatric disorders

We found that positive psychotic experiences in the UK Biobank shared a moderate to substantial amount of common genetic variation with depression and schizophrenia, specifically experiences of auditory and visual hallucinations (un-real voices and un-real vision) and paranoid beliefs (believing in conspiracies) with schizophrenia (r_g_ = 0.27 - 0.67) and with depression (r_g_ = 0.41 - 0.96), consistent with an independent report (55). Our MR analyses indicated that liability to schizophrenia may be directionally associated with a propensity to believe in un-real conspiracies, and that liability to depression may be directionally associated with a propensity to report un-real voices and conspiracies. This latter finding concurs with a twin study showing that depression and psychotic experiences phenotypically influence one another in adolescence over and above genetic influences (56). However, these MR effect sizes were small and not consistently replicated in sensitivity analyses and are therefore unlikely to reflect true causal effects. We also observed large standard errors when using positive psychotic experiences as exposures to predict psychiatric disorders. This may indicate that the UK Biobank GWAS of psychotic experiences was underpowered to detect effects in MR.

This study replicated previous findings of genetic overlap between adolescent PENS and psychiatric disorders (18). In contrast to the findings from the original GWAS, we did not detect a negative genetic association between bipolar disorder and adolescent paranoia and hallucinations and found evidence for genetic overlap between depression and adolescent paranoia and hallucinations. We used more recent summary statistics from larger GWAS of depression and bipolar disorder (40, 41) which likely explain these differences. Our results add further support to previous evidence of genetic associations between adolescent PENS and psychiatric disorders (18, 23).

Schizotypy did not appear to share common additive genetic influences with schizophrenia nor with bipolar disorder, apart from suggestive evidence of genetic overlap between hypomania and depression. There are known phenotypic associations between schizotypy and subsequent risk of broadly defined schizophrenia-spectrum disorders (11), but longitudinal evidence of an association between schizotypy and subsequently being diagnosed with schizophrenia is limited (57) and did not replicate in another study (58). The existing genetic data analysed here do not provide evidence that this association is due to common additive genetic causes.

We found evidence that some genes may be involved in both psychotic experiences and psychiatric disorders. Seven genes were implicated in both positive psychotic experiences in the UK Biobank and psychiatric disorders, including *ANK3*, a gene in the 10q2 1.2 region that has previously been associated with bipolar disorder and schizophrenia (59-61). *TSNARE1* on band 8q24.3, associated with un-real communications and schizophrenia, has previously been implicated in schizophrenia and cognitive function (39, 62). The gene *SNX7* (lp21.3) was mapped to un-real visions, un-real communications and schizophrenia and has known associations with mathematical ability (63). Three neighbouring genes located on band 12q24.11, *FAM216A, PPP1CC* and *GPN3*, and a nearby gene *TMEM116* (12q24.13), overlapped between un-real visions and schizophrenia and have been associated with white blood cell and platelet count, heart rate and body mass index (64-66).

Six mapped genes overlapped between adult schizotypy and psychiatric disorders, including *ANK3* discussed above. As noted elsewhere (20), we found evidence that *CACNA1C* on band 12pl3.33, a gene that has often been associated with psychiatric and developmental disorders (39, 59, 60, 67, 68), may be involved in hypomania, schizophrenia and bipolar disorder. Located within the same region as *CACNA1C* (12p13.33), the genes *DCP1B, TSPAN9* and *FKBP4* also overlapped between hypomania and schizophrenia and have previously been associated with a range of physical health phenotypes including obesity and waist-hip ratio (65, 69). Findings indicated the possible involvement of *CLK1* on 2q33.1 in perceptual aberrations and schizophrenia, a gene previously associated with breast cancer (70).

Our results suggest that psychotic experiences may share more genome-wide genetic overlap with schizophrenia and major depression in adulthood than adolescence. Participants had narrow age ranges in the adolescent PENS (15 - 19 years old) and adult schizotypy (all born in the same year) samples, and were assessed on current psychotic experiences whereas participants in the UK Biobank had a wider and older age range (40 to 69 years old) and were asked to report on psychotic experiences that occurred at any age. It is therefore possible that psychotic experiences reported in the UK Biobank may have been persistent rather than transitory. According to the persistent-impairment model, those with persistent psychotic experiences are at higher risk of developing psychotic disorders and may therefore share more genetic influences with psychiatric disorders (71). It is also possible that some of those in the adolescent samples may be genetically liable to psychotic experiences but will have not yet developed them. A previous study found that genetic liability to schizophrenia was positively associated with adolescent paranoia and hallucinations only after excluding non-zero scores (18), suggesting that genetic liability to schizophrenia may vary in adolescents who did not (yet) report psychotic experiences. This may explain why we found that genetic overlap between psychotic experiences and psychiatric disorders increased with age.

### Stability of genetic influences on psychotic experiences across age

We did not find significant genetic overlap across age and measures for the PENS (adolescence), schizotypy (adulthood) and positive psychotic experiences (adulthood). It is possible that different common additive genetic influences may be involved in psychotic experiences across the lifespan or that the different assessments measured genetically unrelated phenotypes. Another reason why no genetic overlap was found may be the smaller GWAS sample sizes for adolescent PENS and adult schizotypy. It is also possible that non-additive genetic influences or rare variants are shared across psychotic experiences measures and age. In an independent report, genetic risk for psychotic experiences in adults did not predict psychotic experiences in 12 year olds or 16 year olds, in agreement with our findings of lack of genetic overlap across adolescent and adult psychotic experiences, although there was some evidence of genetic association with psychotic experiences at age 18 years (55). Because UK Biobank participants were asked to report on psychotic experiences across their lives, some may have referred to experiences such as hallucinations in later adulthood that can occur as part of conditions such as dementia, eye conditions, Charles Bonnet syndrome and medication side effects, which may have different etiologies to psychotic experiences in adolescence.

The gene association and mapping analyses suggested that some of the same genes may be involved in psychotic experiences across age. The gene *ANK3* (discussed above) was implicated for adult schizotypy (physical anhedonia) and positive psychotic experiences (un-real visions). The gene *PAN3* on band 13q12.2 was implicated in adolescent PENS (cognitive disorganization) and adult schizotypy (perceptual aberrations). *NADK2* (mitochondrial) on 5p13.2 was mapped to both negative symptoms in adolescence and to positive psychotic experiences during adulthood (unreal communications).

### Within-sample associations between specific types of psychotic experiences

We also investigated within-sample genetic associations and found that un-real voices and un-real visions shared the same common genetic influences in adults and were bidirectionally associated in Mendelian Randomization analysis. This bidirectional association may indicate a pervasive shared genetic basis between auditory and visual hallucinatory experiences rather than true causal associations because pleiotropic effects of the genetic instruments through a latent common precursor, and therefore violations of the MR assumptions, seem likely. As such, experiences resembling auditory and visual hallucinations may share biological pathways. Our findings of genetic correlations between the three psychiatric disorders we tested are in line with previous findings (26, 41), while we used more recent GWAS results for bipolar disorder.

### Limitations

We note caution in interpreting MR results that compared within-sample psychotic experiences because sample overlap could, in within-sample MR, lead to weak instrument bias away from the null (72). Weak instrument bias may also have affected analyses that used psychotic experiences as the exposure, where genetic variants below conventional genome-wide significance levels were used as instruments. In two-sample MR, weak instruments create bias toward the null and as a result true causal effects may not have been detected. Future studies could re-evaluate these causal associations once larger GWAS for psychotic experiences become available.

Many of the overlapping genes we report were identified based only on chromatin interactions. The chromatin interaction data had a high resolution (10kb) resulting in more mapped genes, and included enhancer-promoter and promoter-promoter interactions only (73), therefore providing strong hypotheses for variant-gene associations. For psychotic experiences and depression, functional consequences on genes were annotated to variants clumped at lower than conventional genome-wide significance levels. As such, these results will benefit from replication once better powered GWAS become available.

### Implications and conclusions

Positive psychotic experiences during adulthood and PENS during adolescence share genetic influences with psychiatric disorders. This study implicated specific genes that may be involved in both psychotic experiences in the community and psychiatric disorders, and genes associated with psychotic experiences across development. Future molecular genetic studies could further investigate whether these genes may be involved in biological pathways relevant to a broader psychosis spectrum. Subclinical experiences of auditory and visual hallucinations in adults may have similar biological etiologies. Our findings and other independent reports indicate that psychotic experiences assessed using different measures at different developmental stages may not reflect genetically similar phenomena.

## Acknowledgements

This work was supported by the UK Medical Research Council (G1100559 to AR) and a Wellcome Trust ISSF grant to AR. WB is funded by the Camara-Rijvers David Studentship.

The authors gratefully acknowledge the ongoing contribution of the participants in the Twins Early Development Study (TEDS) and their families, the Avon Longitudinal Study of Children and Parents (ALSPAC), the Child and Adolescent Twin Study in Sweden (CATSS) and those who participated in the North Finland Birth Cohort (NFBC) and in the UK Biobank. We thank the funding bodies and research teams which includes interviewers, computer and laboratory technicians, clerical workers, research scientists, volunteers, managers, receptionists and nurses. TEDS is supported by a program grant to Robert Plomin from the UK Medical Research Council (MR/M021475/1). The UK Medical Research Council and Wellcome (Grant ref: 102215/2/13/2) and the University of Bristol provide core support for ALSPAC. This publication is the work of the authors and WB and AR will serve as guarantors for the contents of this paper. ALSPAC GWAS data was generated by Sample Logistics and Genotyping Facilities at Wellcome Sanger Institute and LabCorp (Laboratory Corporation of America) using support from 23andMe. We are grateful to Neale Lab and the Psychiatric Genomics Consortium (PGC) for their contributions to providing the genetic summary results used in this study and to Dr. W. Hennah and A. Ortega-Alonso for preparing and sharing the schizotypy summary statistics.

