## Supplementary Figures and Supplementary Note for "Genetic overlap between psychotic experiences in the community across age and with psychiatric disorders"

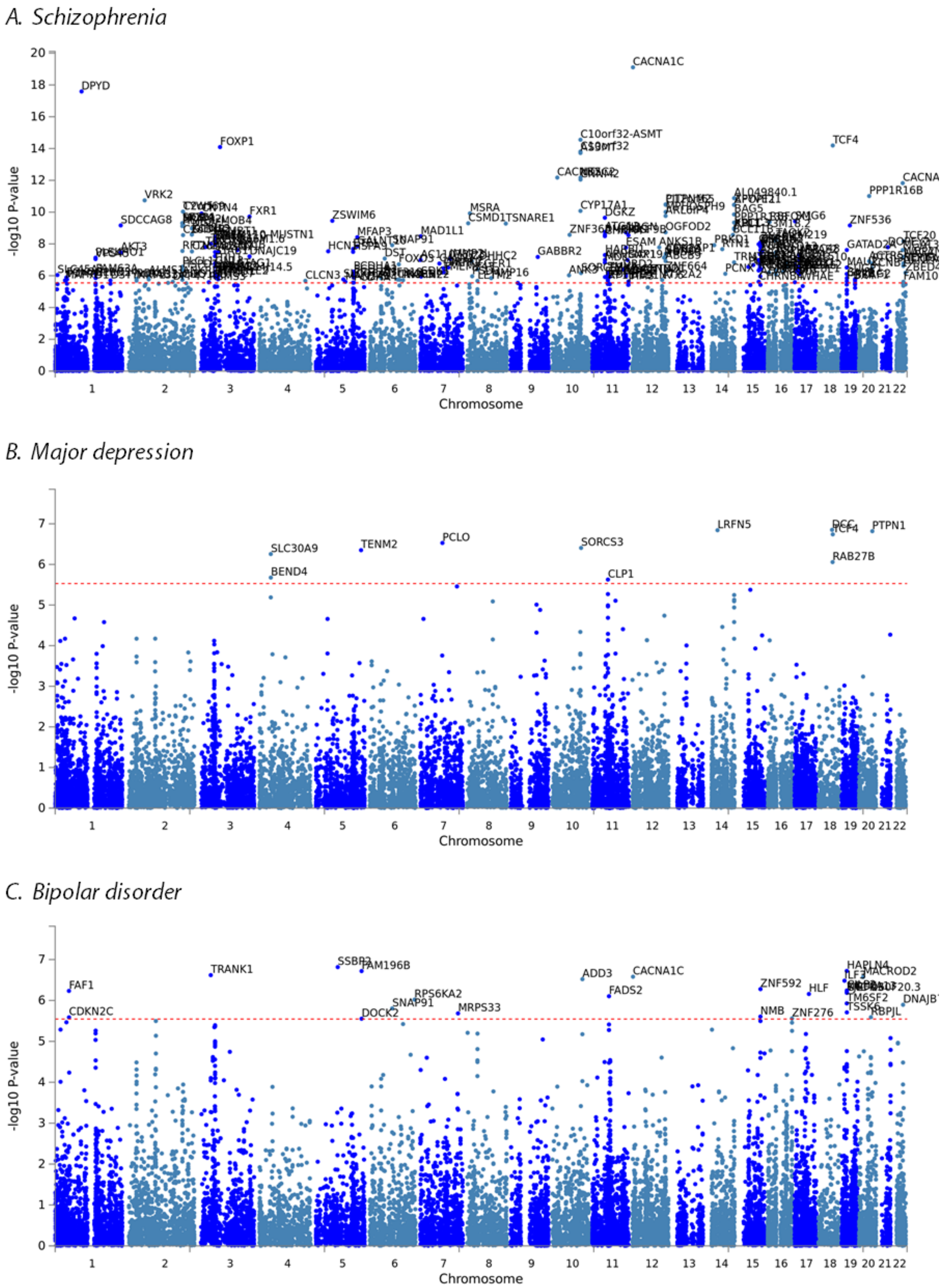

***Figure S1.*** **Manhattan plots for MAGMA gene-based analyses.**

*Note:* Red dashed line indicates genome-wide significance for schizophrenia as defined at p = 0.05/17674 genes tested = 2.83 x $10^{-6}$, for major depression at p = 0.05/16943 = 2.95 x $10^{-6}$ and for bipolar disorder at p = 0.05/17538 = 2.85 x $10^{-6}$.

*D. Un-real voice*

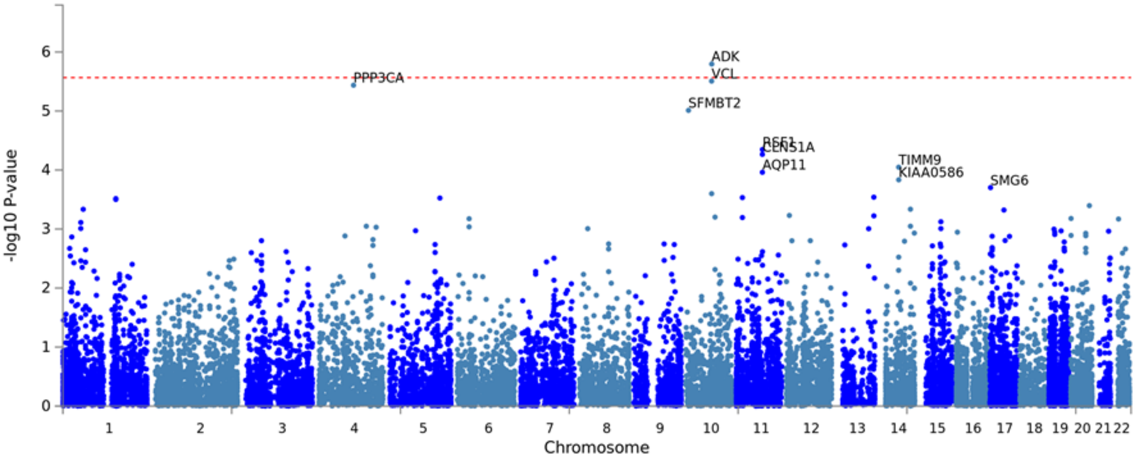

*E. Un-real visions*

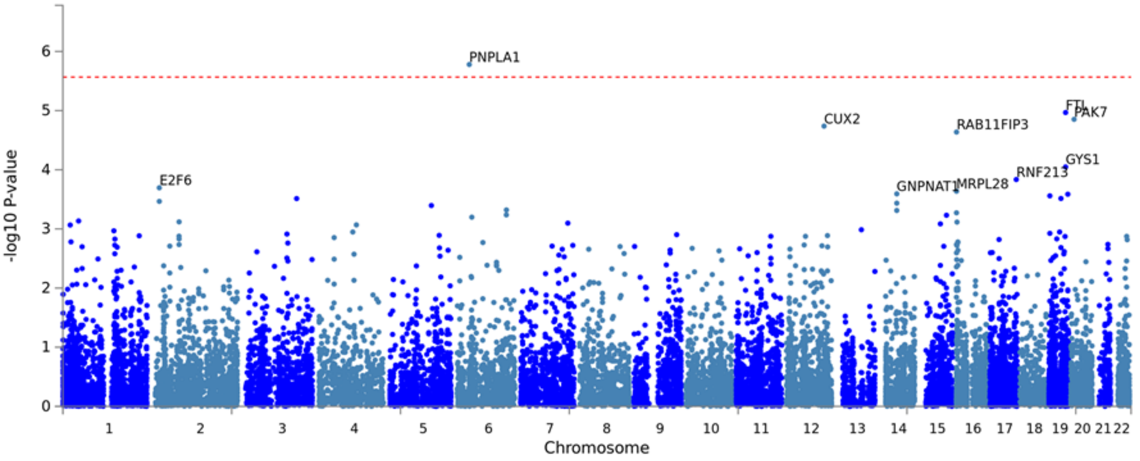

*F. Un-real conspiracies*

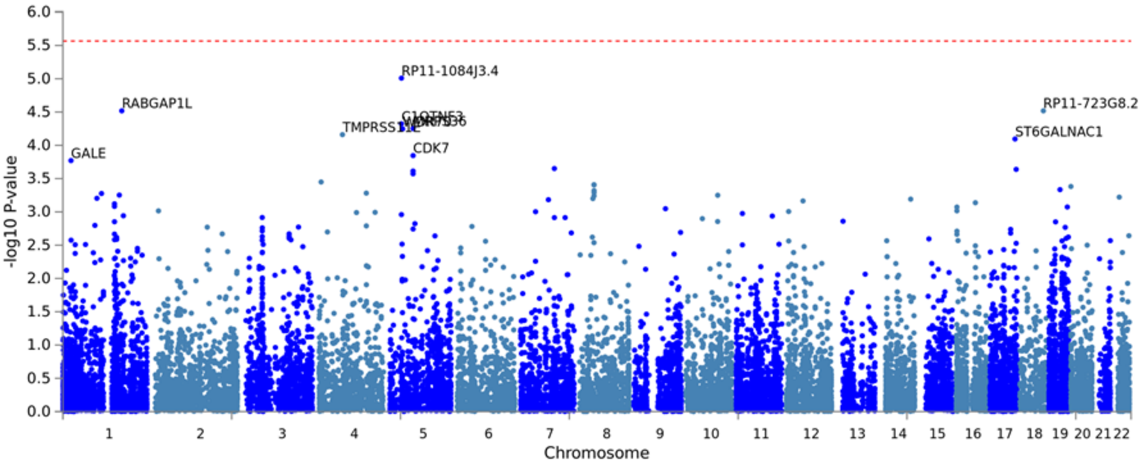

***Figure S1.*** **Manhattan plots for MAGMA gene-based analyses (continued).**

*Note:* Red dashed line indicates genome-wide significance for positive psychotic experiences in the UK Biobank as defined at $p = 0.05/18423$ genes tested $= 2.71 \times 10^{-6}$.

*G. Un-real communications*

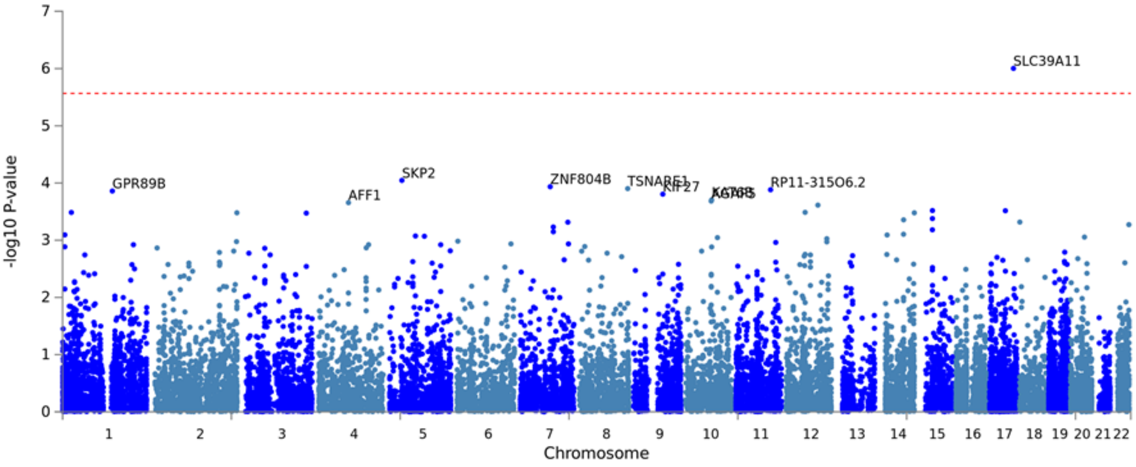

*H. Hypomania*

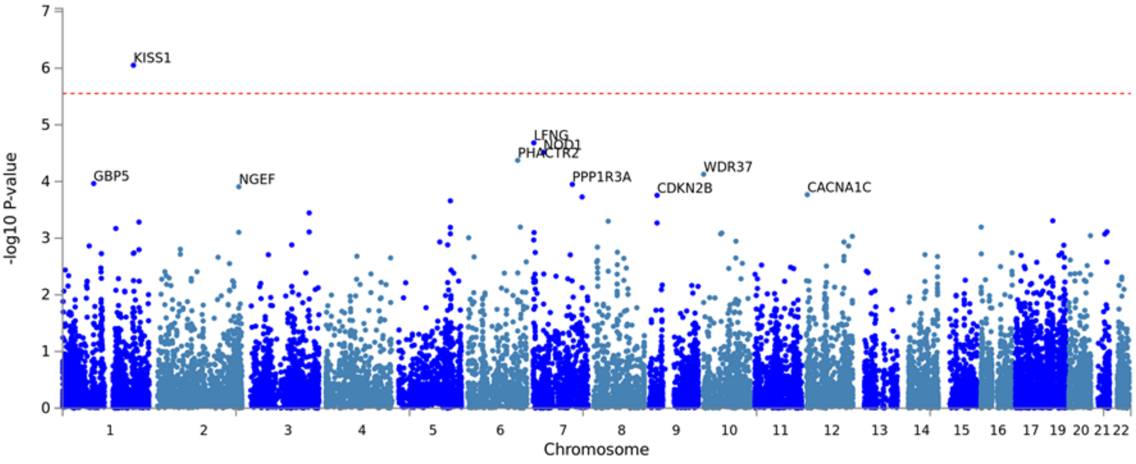

*I. Perceptual Aberrations*

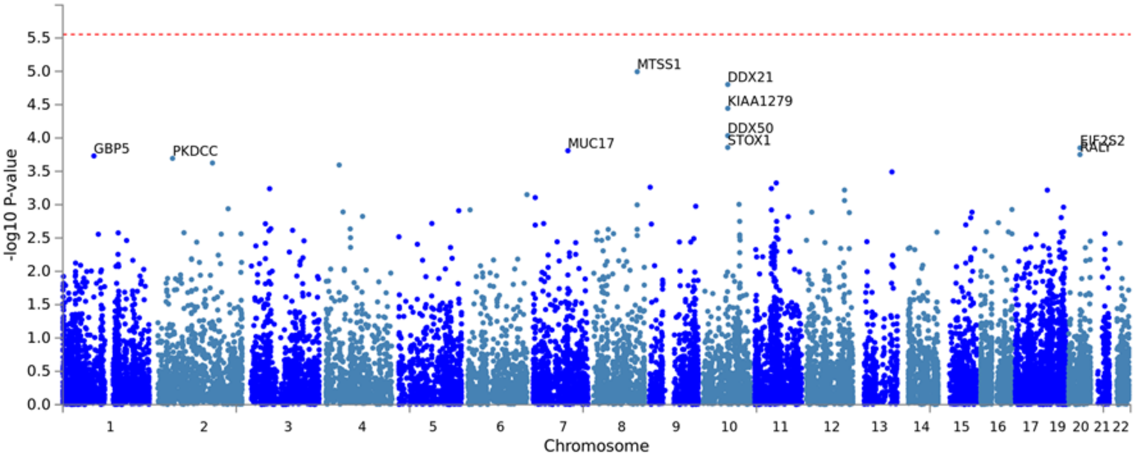

***Figure S1.*** **Manhattan plots for MAGMA gene-based analyses (continued).**

Note: Red dashed line indicates genome-wide significance for positive psychotic experiences in the UK Biobank at defined at P = 0.05/18423 genes tested = 2.714e-6 and for schizotypy in the North Finland Birth Cohort at P = 0.05/17895 = 2.794e-6.

*J. Physical anhedonia*

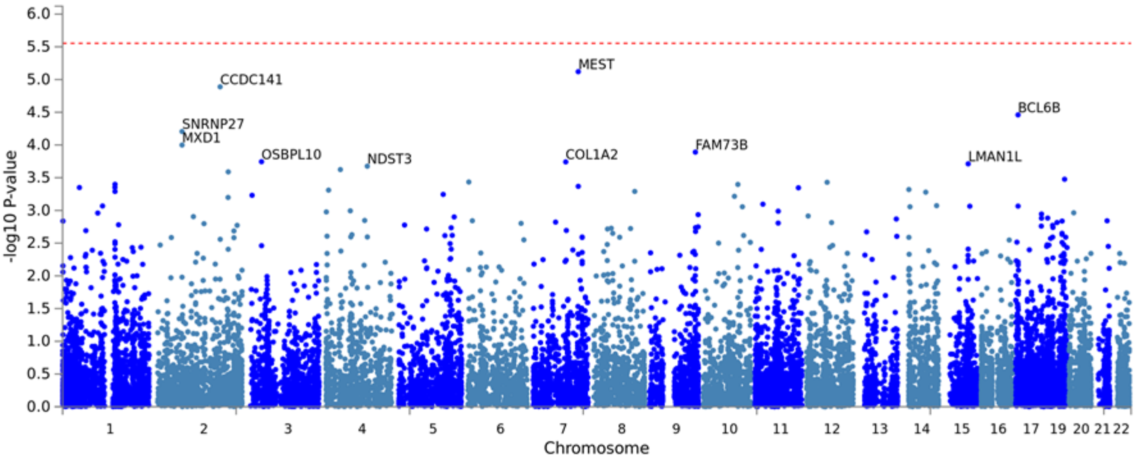

*K. Social anhedonia*

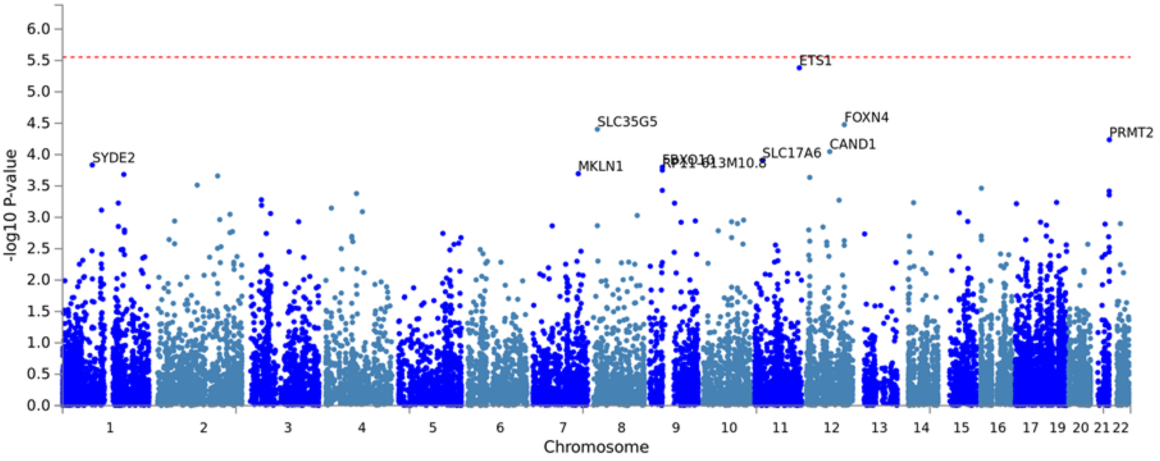

*L. Paranoia and hallucinations*

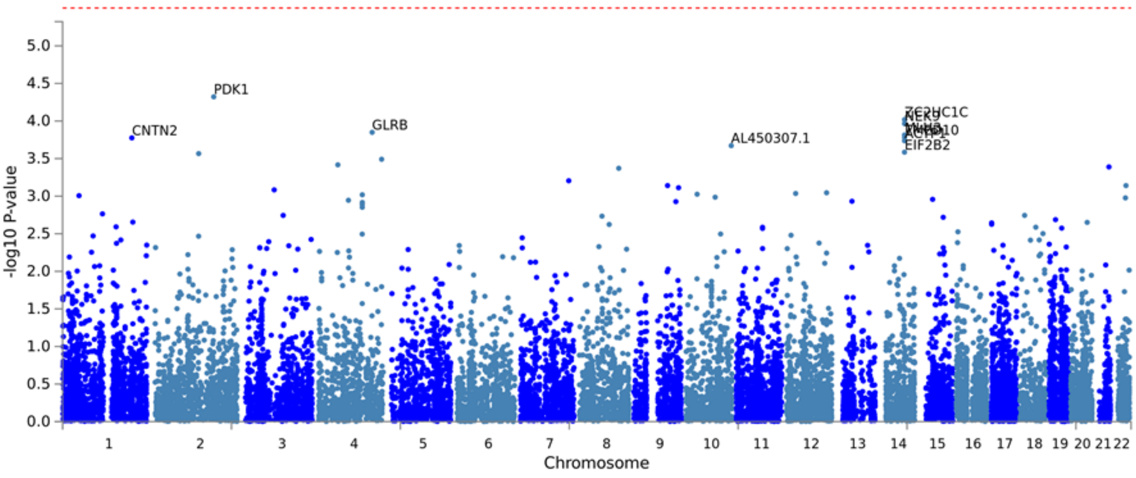

***Figure S1.*** **Manhattan plots for MAGMA gene-based analyses (continued).**

Note: Red dashed line indicates genome-wide significance for schizotypy in the North Finland Birth Cohort at p = 0.05/17895 genes tested = 2.794e-6 and for adolescent psychotic experiences and negative symptom traits at p = 0.05/15957 = 3.133e-6.

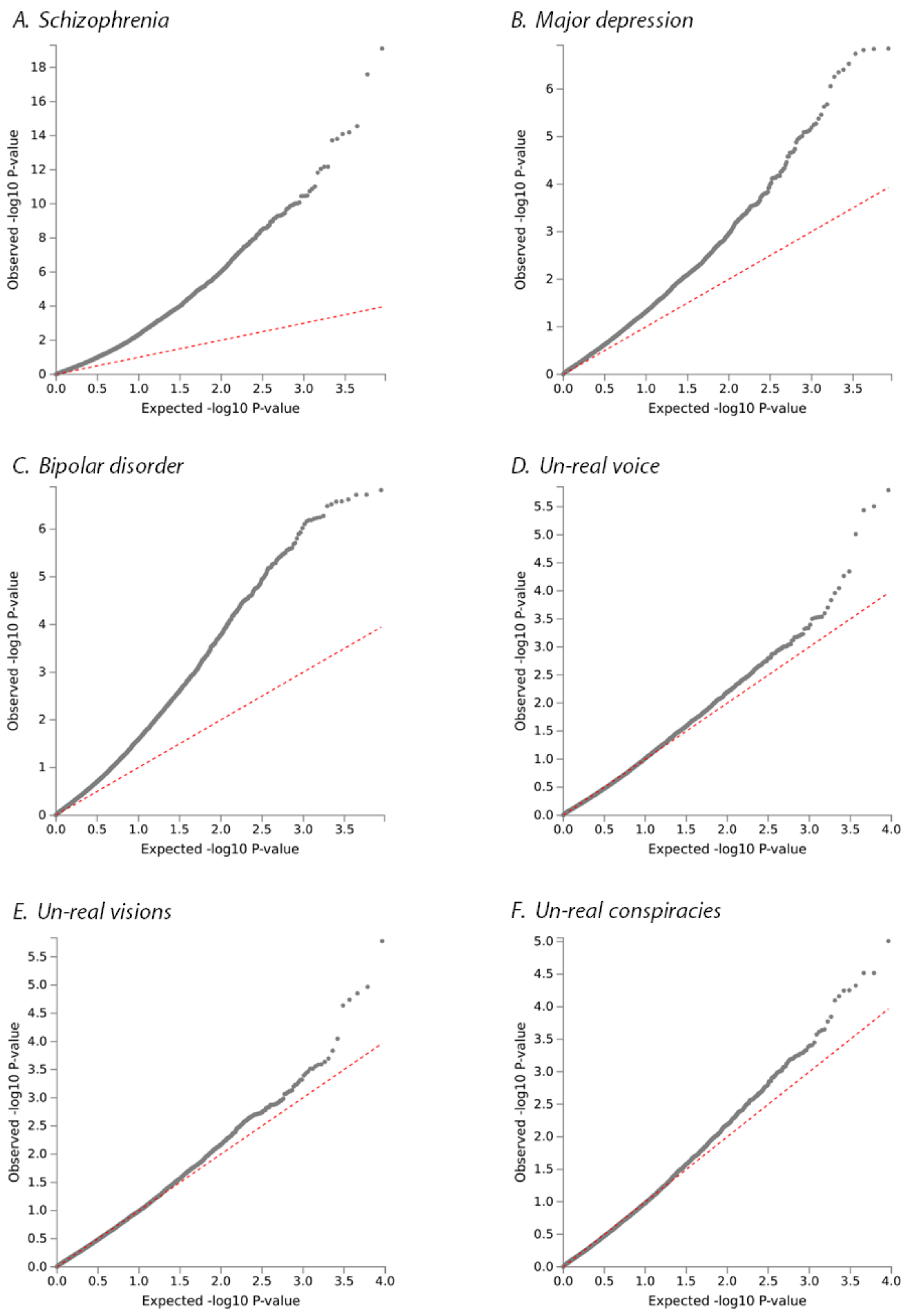

***Figure S2.*** **QQ plots for MAGMA gene-based analyses.**

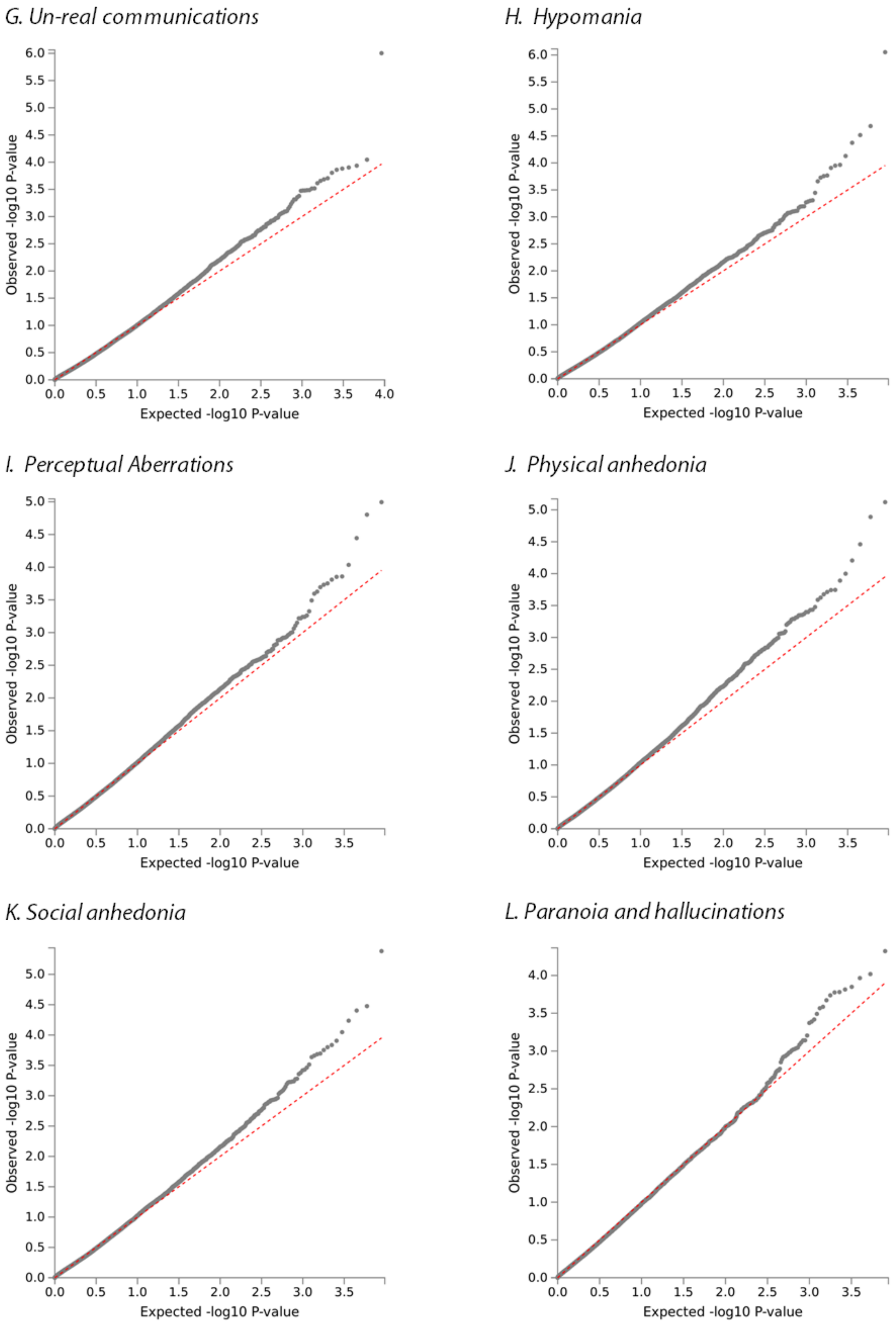

***Figure S2.*** **QQ plots for MAGMA gene-based analyses (continued).**

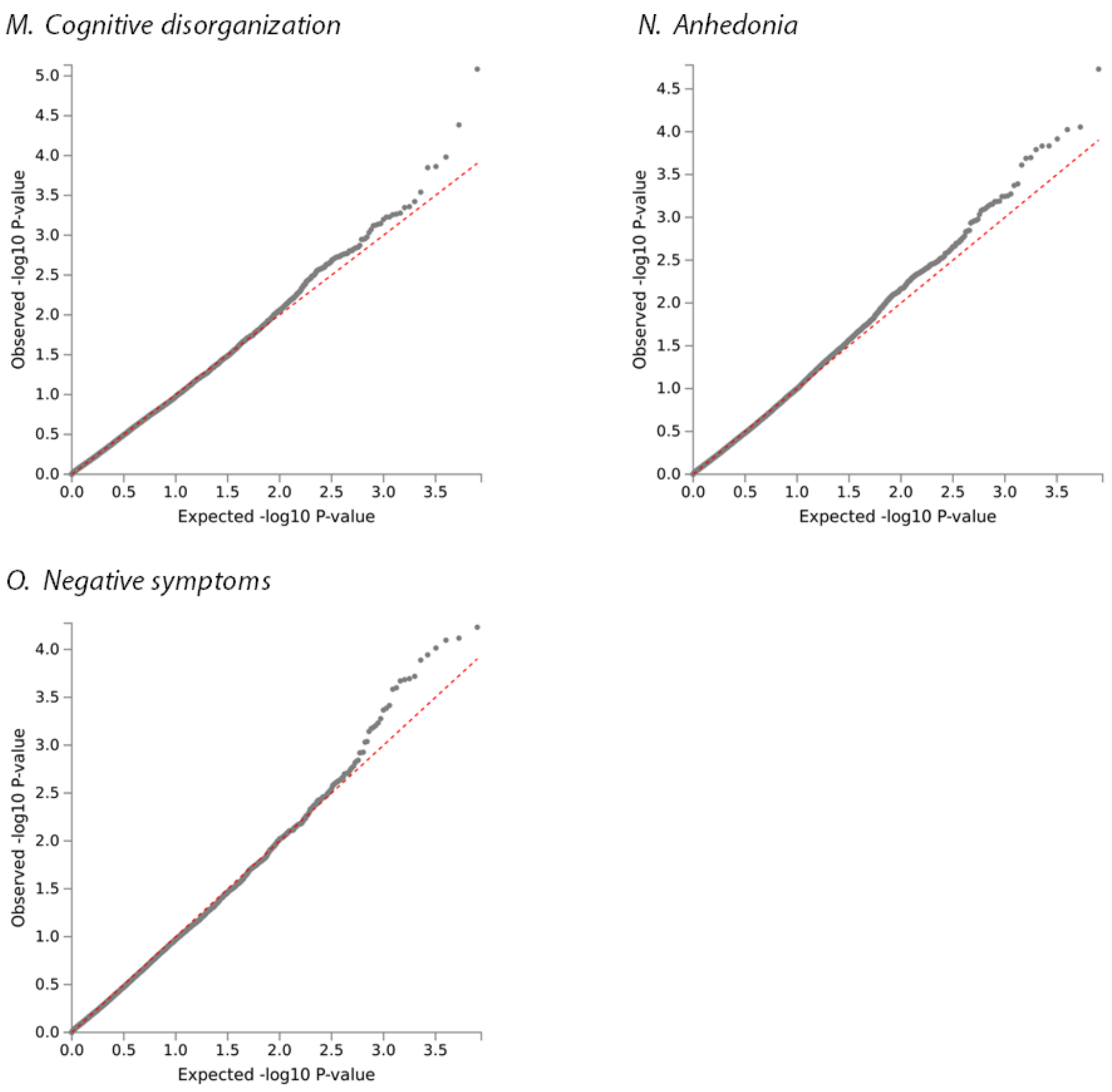

***Figure S2.*** **QQ plots for MAGMA gene-based analyses (continued).**

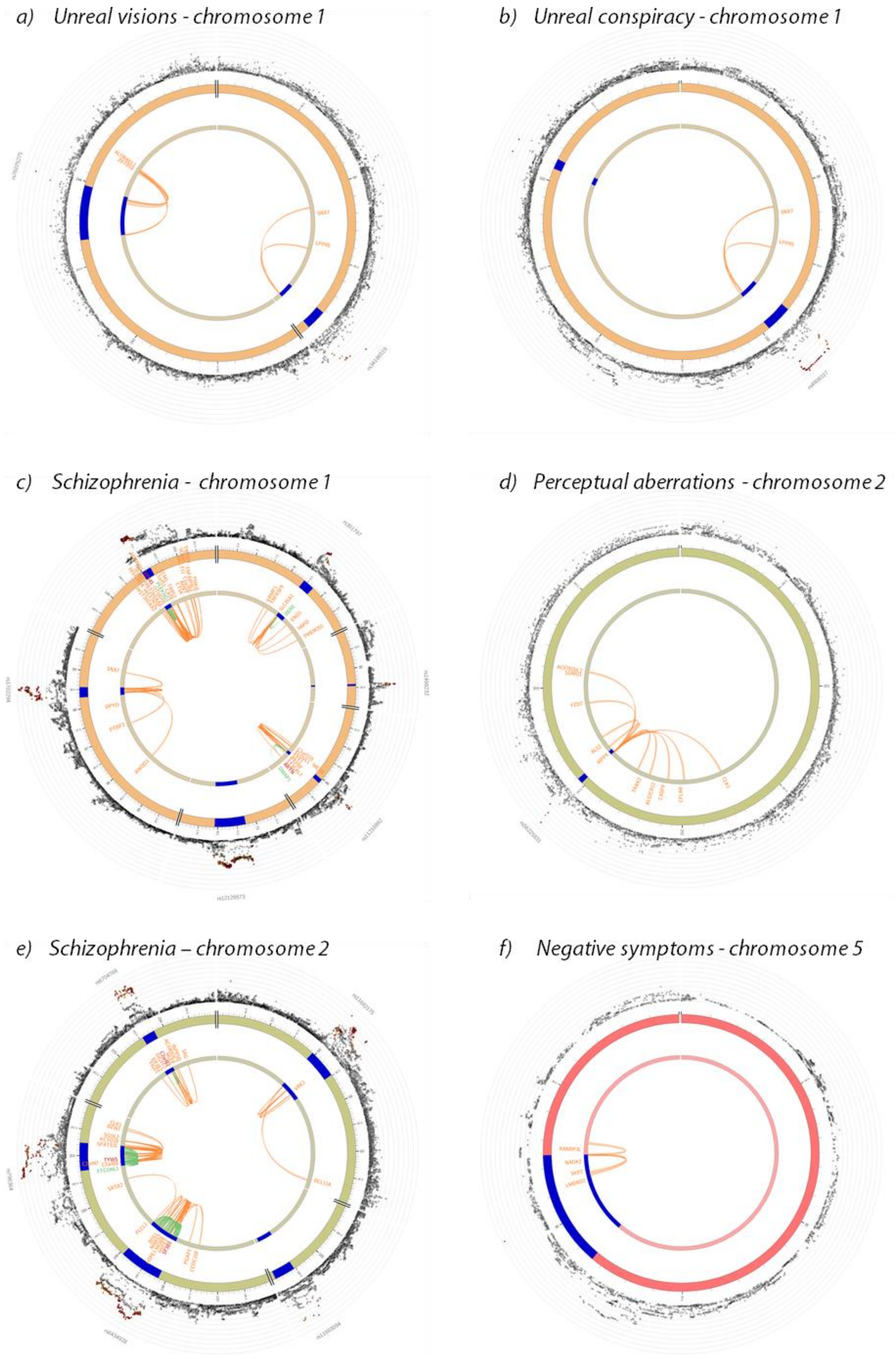

***Figure S3*. Circos plots for chromatin interactions and eQTL.**

*g) Unreal communications - chromosome 5*

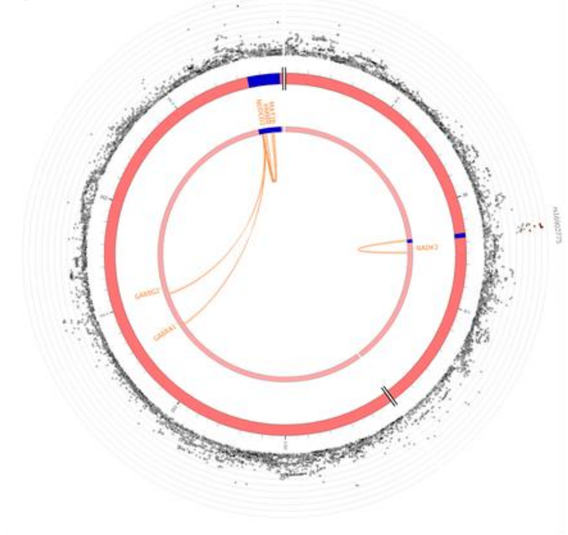

*h) Unreal communications – chromosome 8*

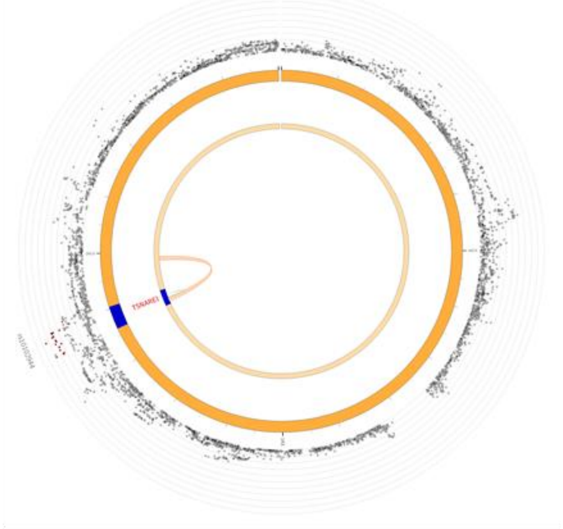

*i) Schizophrenia - chromosome 8*

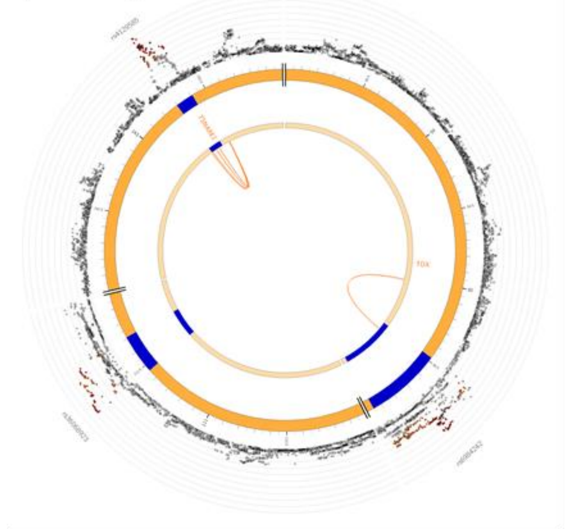

*j) Physical anhedonia - chromosome 10*

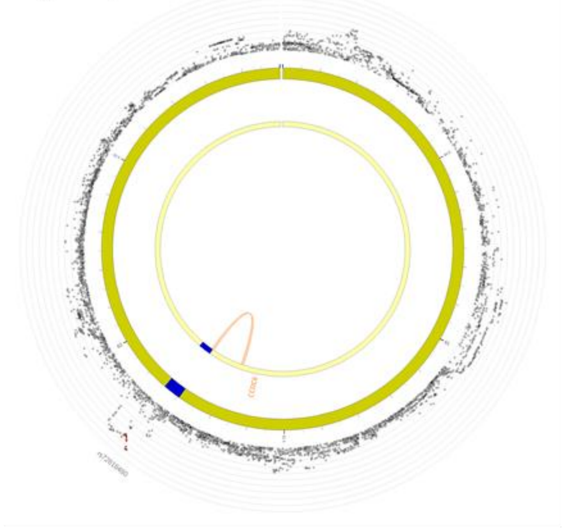

*k) Unreal visions - chromosome 10*

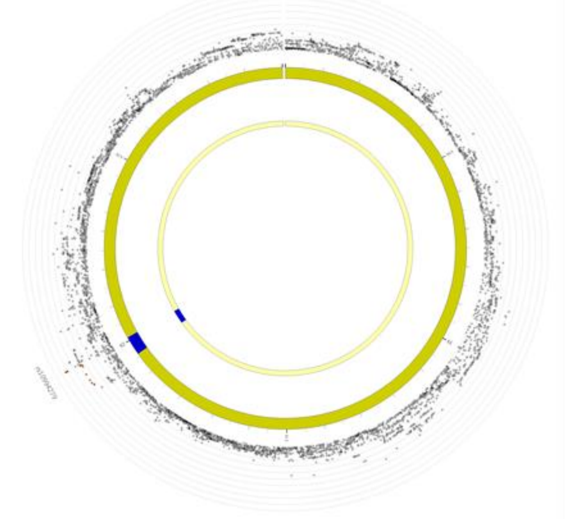

*l) Hypomania - chromosome 12*

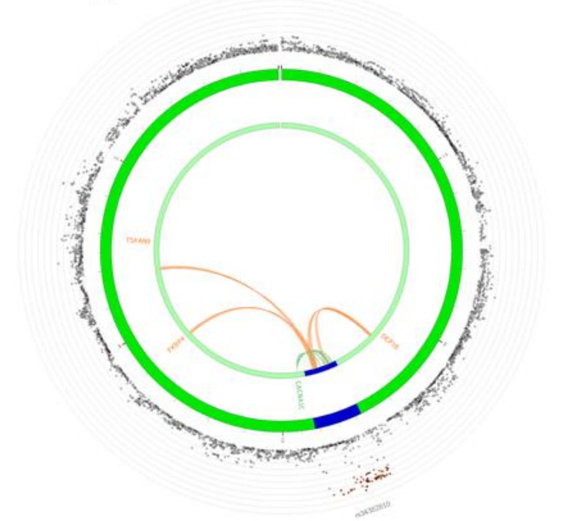

***Figure S3 (continued)*. Circos plots for chromatin interactions and eQTL.**

*m) Unreal visions - chromosome 12*

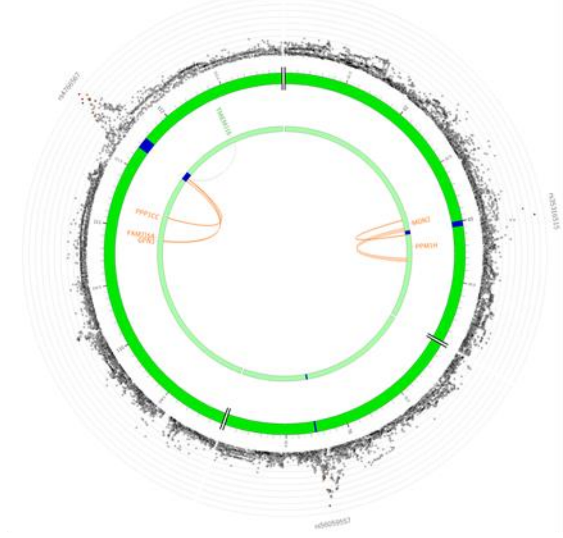

*n) Schizophrenia - chromosome 12*

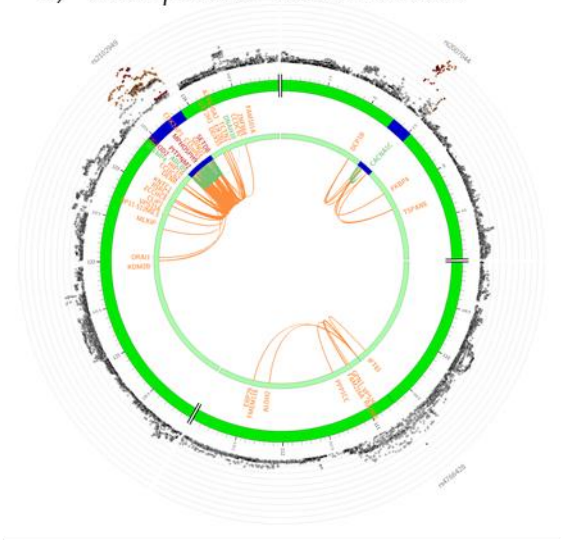

*o) Perceptual aberrations - chromosome 13*

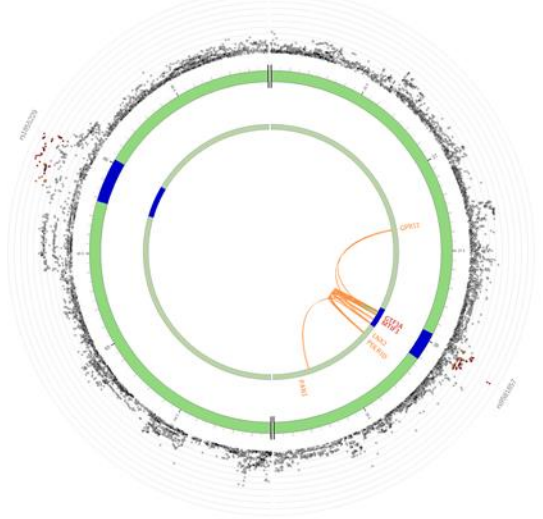

***Figure S3 (continued)*. Circos plots for chromatin interactions and eQTL.**

*Note:* Manhattan plot displayed in outer most ring with loci colour coded according to the amount of LD shared with lead independent SNPs as follows: red ($r^2 > 0.8$), orange ($r^2 > 0.6$), green ($r^2 > 0.4$) and blue ($r^2 > 0.2$). Genomic risk loci are displayed in blue on the chromosome ring (second and third layers). Genes mapped by chromatin interaction are displayed in orange, by eQTLs in green, and by both chromatin interaction and eQTLs in red; Mapped genes that overlapped between phenotypes are listed in Table 3.

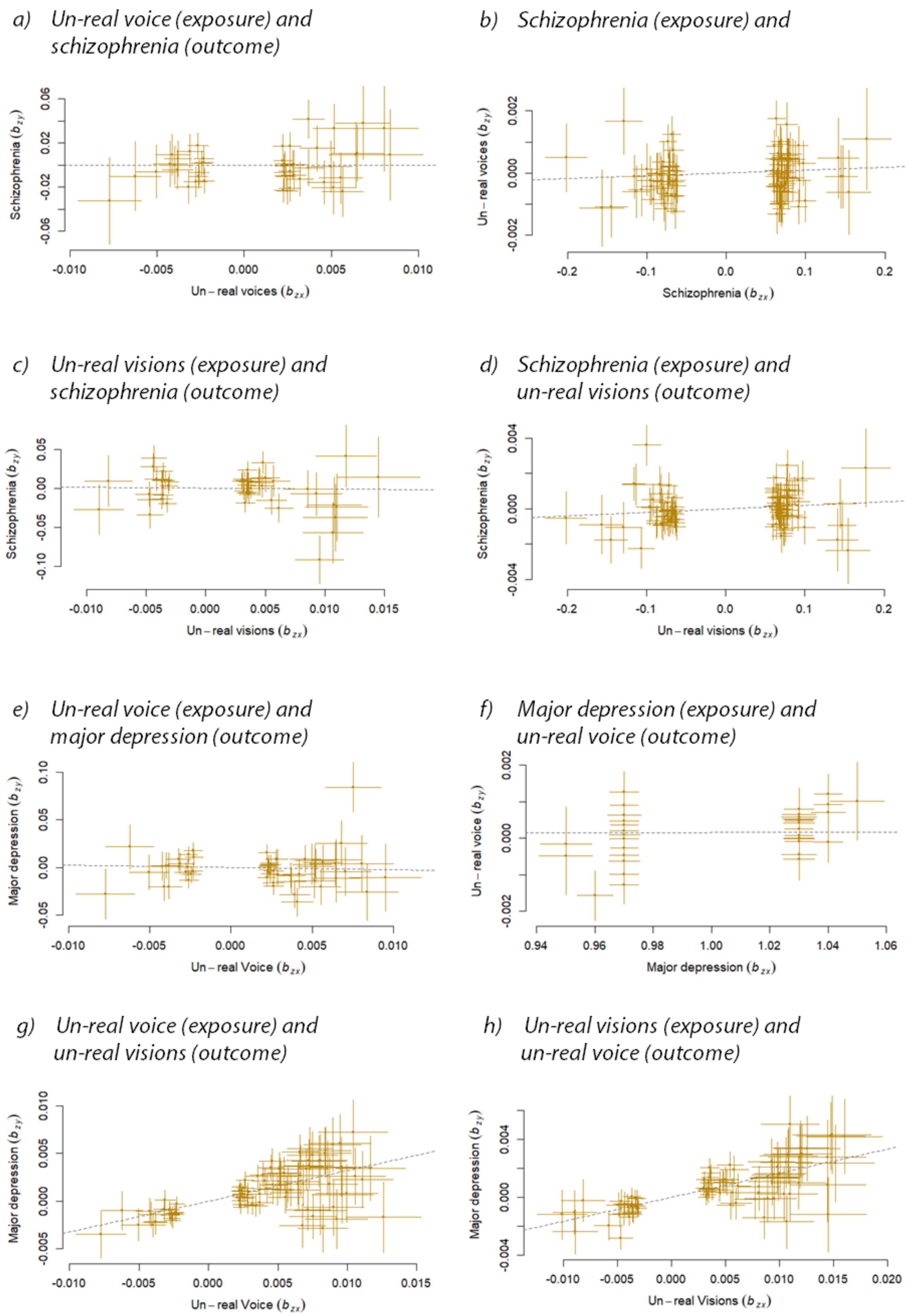

***Figure S4*. Generalised Summary-Based Mendelian Randomisation analyses.**

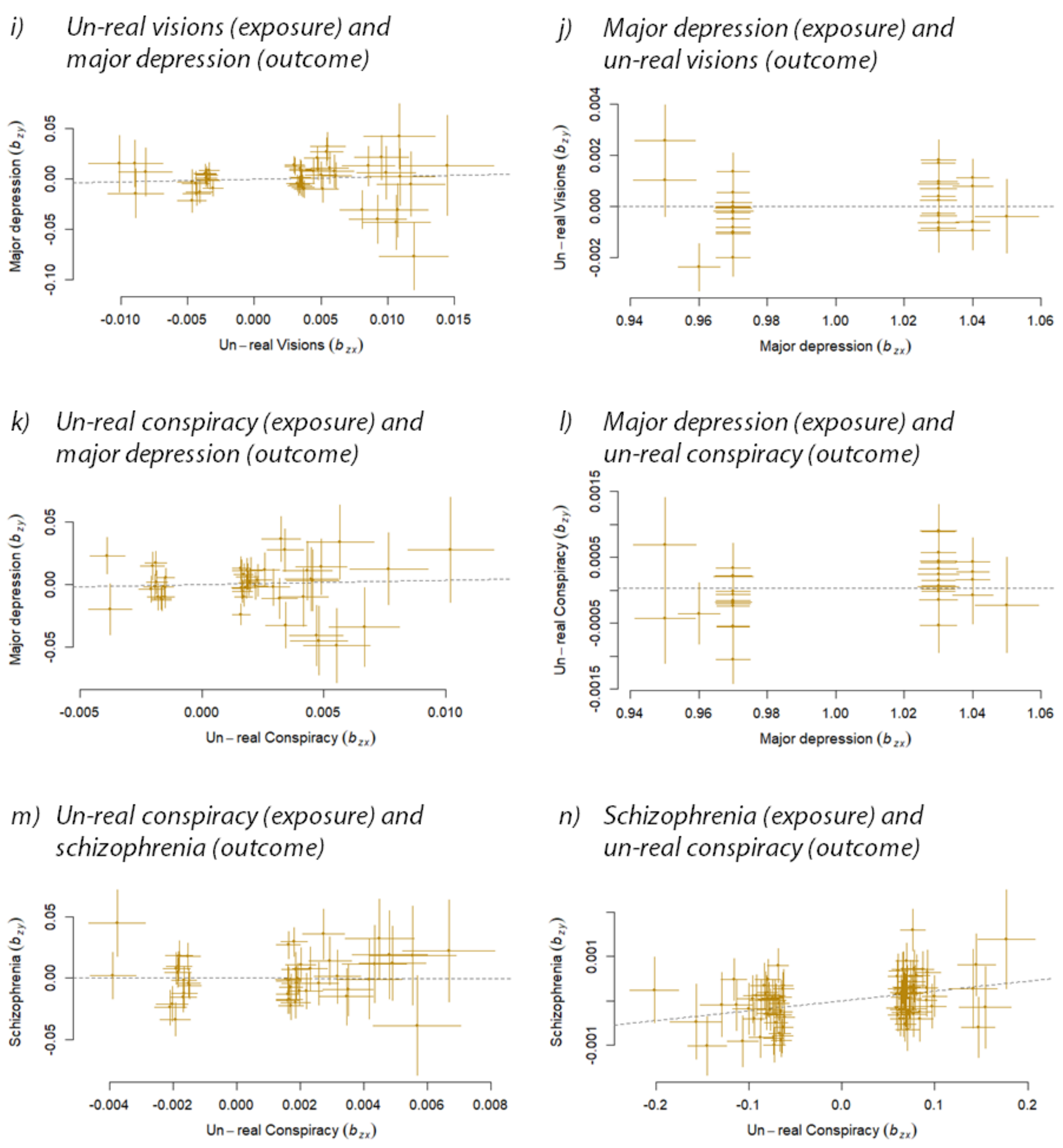

***Figure S4.*** **Generalised Summary-Based Mendelian Randomisation analyses (continued).**

*Note:* Scatterplots with the x-axis displaying instrumental variable effects on the exposure ($b_{zx}$) and the y-axis displaying the instrument-outcome association ($b_{zy}$). Regression lines included for reference.

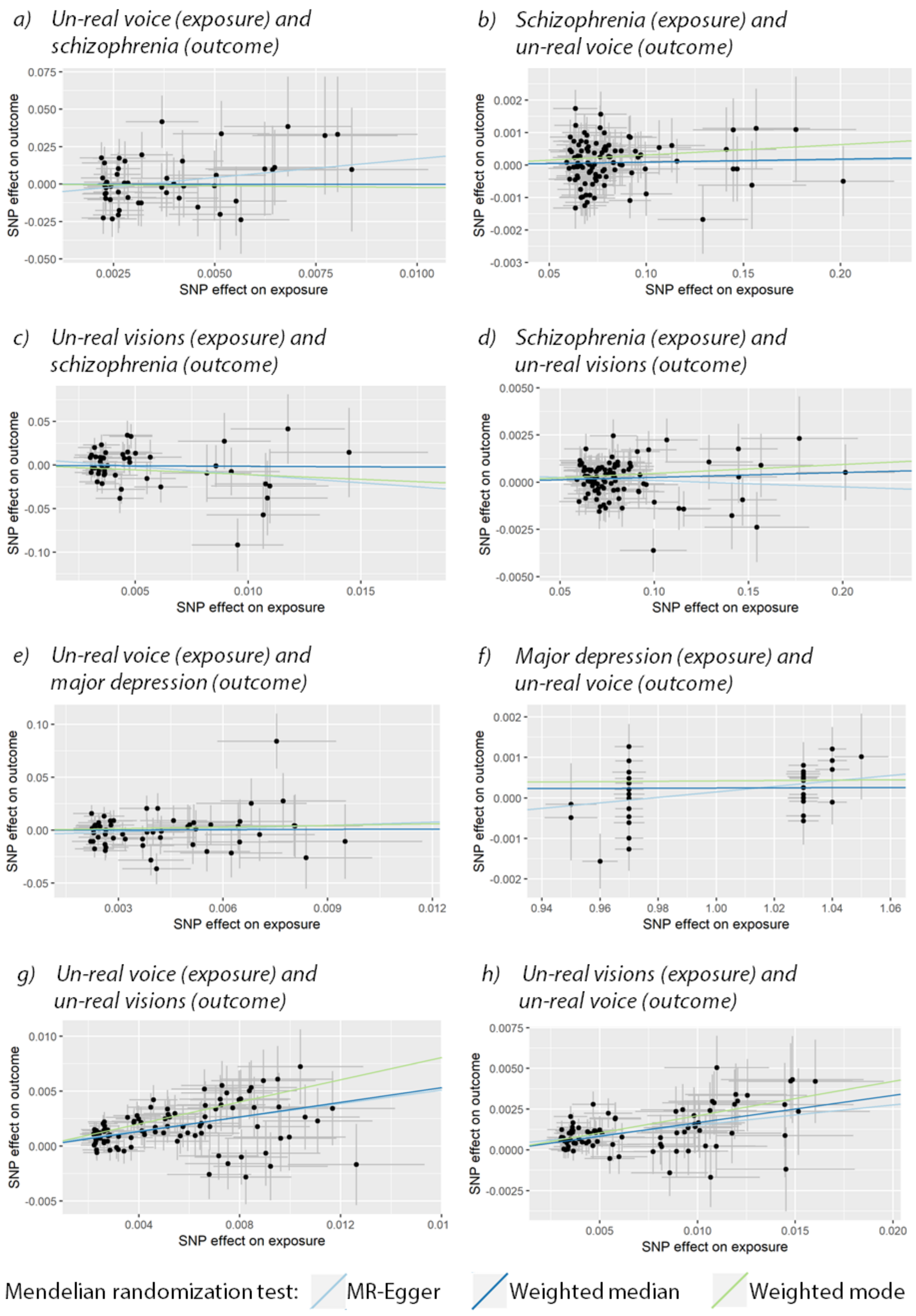

***Figure S5.*** **MR-Egger, Weighted Median and Weighted Mode Mendelian randomisation sensitivity analyses.**

*i)* *Un-real visions (exposure) and major depression (outcome)*

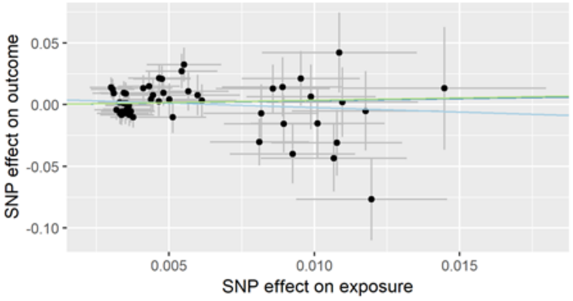

*j)* *Major depression (exposure) and un-real visions (outcome)*

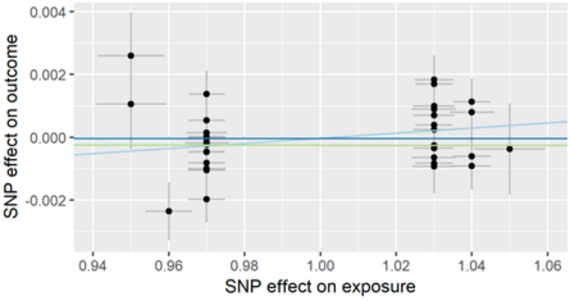

*k)* *Un-real conspiracy (exposure) and major depression (outcome)*

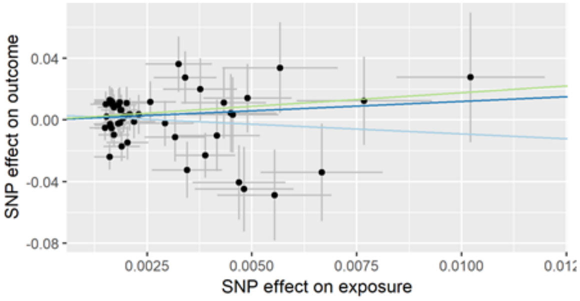

*l)* *Major depression (exposure) and un-real conspiracy (outcome)*

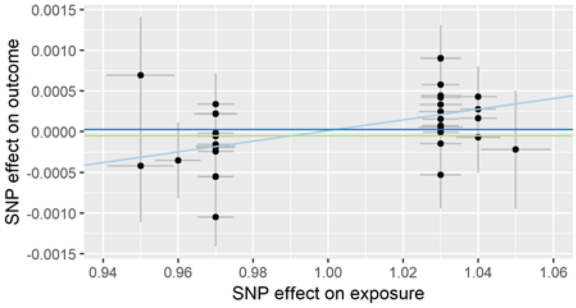

*m)* *Un-real conspiracy (exposure) and schizophrenia (outcome)*

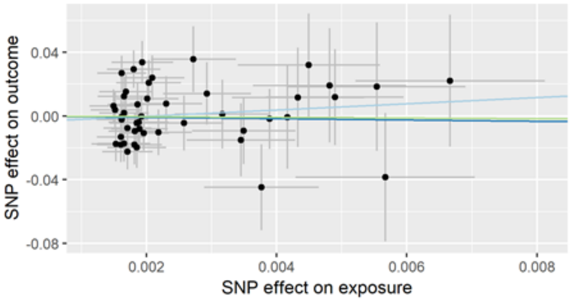

*n)* *Schizophrenia (exposure) and un-real conspiracy (outcome)*

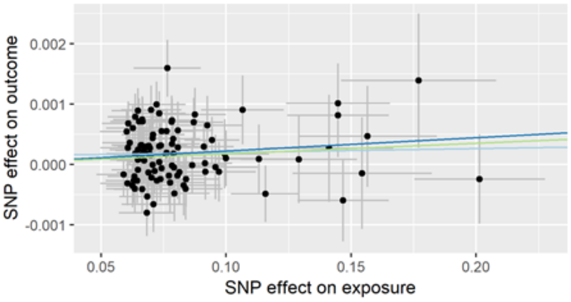

***Figure S5.*** **MR-Egger, Weighted Median and Weighted Mode Mendelian randomisation sensitivity analyses (continued).**

### Extended methods

*Adolescent psychotic experiences and negative symptom traits*

The ALSPAC sample (1, 2) invited pregnant women resident in Avon, UK and with an expected delivery date between 1st April 1991 and 31st December 1992 to participate in the study. The initial sample consisted of 14,775 children. Informed consent for the use of data collected via questionnaires and clinics was obtained from participants following the recommendations of the ALSPAC Ethics and Law Committee at the time. Consent for biological samples has been collected in accordance with the Human Tissue Act (2004). Please note that the ALSPAC study website contains details of all the data that is available through a fully searchable data dictionary and variable search tool (http://www.bristol.ac.uk/alspac/researchers/our-data/).

Ethical approval for the original adolescent PENS GWAS (3) was obtained for ALSPAC from the ALSPAC Ethics and Law Committee and the Local Research Ethics Committees, for TEDS from the Institute of Psychiatry ethics committee (ref: 05/Q0706/228), and for CATSS from the Karolinska Institute Ethical Review Board.

The harmonisation process of items for psychotic experiences and negative symptom traits (PENS) across TEDS, CATSS and ALSPAC was informed by principle component analyses, an expert clinical team and the availability of overlapping items (3). For instance, paranoia and hallucinations items from SPEQ that were harmonised across the three adolescent PENS cohorts were "*How often have you thought 'I might be being observed or followed?'*" , for anhedonia "*When something exciting is coming up in my life, I really look forward to it"* (reverse scored), for cognitive disorganisation "*Do you find it difficult in controlling your thoughts?*" and for parent-rated negative symptoms "*My child has a lack of energy and motivation*". Items for anhedonia was not available in the CATSS sample and for cognitive disorganisation unavailable in ALSPAC.

Linear regression GWAS and Generalized Estimating Equation (GEE; to account for the presence of twin pairs) was performed on the four PENS scales (3). Summary result files were obtained from the authors with permission from the original study cohorts.

*Schizotypy during middle adulthood*

Four schizotypy scales were used to assess psychotic experiences during middle adulthood: Perceptual aberrations were assessed with the Perceptual Aberration Scale (4) with 35 true/false items devised to assess experiences in the general population that resemble clinical features of schizophrenia with an emphasis on body image aberrations including unclear body boundaries, body size and physical attributes being distorted, or feelings of estrangement from one's own body. Items also assessed unusual visual and auditory experiences, for example "*My hearing is sometimes so sensitive that ordinary sounds become uncomfortable*" and "*Sometimes when I look at things like tables and chairs, they seem strange*".

Hypomania was from the Hypomanic Personality Scale (5) and consisted of 48 true/false items devised to assess hypomania, gregariousness, grandiosity and euphoria (e.g. "*I can usually slow myself down when I want to"* and "*I have often been so excited about an involving project that I didn't care about eating or sleeping*").

Two scales from Chapman's Schizotypia Scales were employed to assess social anhedonia with the Revised Social Anhedonia Scale and physical anhedonia from the Revised Physical Anhedonia Scale (6), devised to assess the inability to take pleasure from physical (61 true/false items, e.g. "*One food tastes as good as another to me*") and social (40 true/false items, e.g. *"I prefer watching television to going out with other people"*) stimuli respectively.

Summary statistics from linear regression GWAS performed on these four schizotypy scales were obtained from the authors (7).

*Positive psychotic experiences assessed in adults*

GWAS on four dichotomous items from the UK Biobank were included. The items assessed psychotic experiences in adults aged 40-69 years: Whether participants ever experienced hearing an un-real voice (UK Biobank phenotype ID = 20463; *"Did you ever hear things that other people said did not exist, like strange voices coming from inside your head talking to you or about you, or voices coming out of the air when there was no one around?"*), had an un-real vision (UK Biobank phenotype ID = 20471; *"Did you ever see something that wasn't really there that other people could not see?"*), held a belief in an un-real conspiracy (UK Biobank phenotype ID = 20468; *"Did you ever believe that there was an unjust plot going on to harm you or to have people follow you, and which your family and friends did not believe existed?"*) and experienced un-real communications or signs (UK Biobank phenotype ID = 20474; *"Did you ever believe that a strange force was trying to communicate directly with you by sending special signs or signals that you could understand but that no one else could understand (for example through the radio or television)?"*). The mean age of onset of positive psychotic experiences reported by UK Biobank participants was 31.6 (s.d. = 17.6) years. Of those reporting positive psychotic experiences, 11.3% indicated that they have received medication for psychotic experiences and 21.3% have talked to a mental health professional about their psychotic experiences.

Separate linear regression GWAS was performed on each of the four positive psychotic experiences items for individuals of European ancestry (N = 116,787 - 117,794) by Neale Lab and summary results was downloaded from the Neale Lab website (http://www.nealelab.is/uk-biobank).

*Mendelian randomization*

Mendelian randomization (MR)(8) was conducted to further explore the relationship between psychotic experiences and psychiatric disorders for the phenotype pairs that had significant genetic correlations. MR is used to test for a causal relationship between an exposure (the putatively causal trait) and outcome trait by using instrumental variables as proxies for the exposure trait. In MR, instrumental variables are SNPs robustly associated with the exposure based on GWAS results. Due to the random nature of Mendelian segregation of genetic variants during meiosis, the extent to which unmeasured confounding factors influence the outcome is not expected to differ between those who inherited a specific copy of a genetic variant and those who did not (akin to the randomization process employed in randomized controlled trials).

The presence of a causal association between the exposure (X) on the outcome (Y) trait can be calculated as the ratio of the effect size of a SNP instrumental variable (Z) on the outcome over its effect on the exposure: $\hat{b}_{XY} = \hat{b}_{ZY}/\hat{b}_{ZX}$, where $\hat{b}_{XY}$ is the effect of the exposure on the outcome, $\hat{b}_{ZY}$ is the effect of the SNP instrument variable on the outcome and $\hat{b}_{ZX}$ is its effect on the exposure. To overcome the small effect sizes of individual SNPs, multiple SNPs are used as instrumental variables to increase power and an aggregate $\hat{b}_{XY}$ effect can be obtained using a generalized least squares approach (9).

The effect alleles for summary statistics for positive psychotic experiences, schizophrenia and major depression were harmonised to be in phase with the 1000 Genomes (phase 3) reference panel in both the outcome and exposure data. The effects were log odds ratios for binary traits, except for the UK Biobank psychotic experiences for which linear regression coefficients were provided.

*FUMA*

FUMA is a web application that offers a streamlined pipeline to perform several post-GWAS analyses on summary statistics. Summary statistics are uploaded to a server and automatically deleted once the analyses have been performed. Results are stored on the FUMA servers until users remove these.

Post-GWAS functional annotation analyses for adolescent PENS and adult schizotypy was reported in the original GWAS publications (3, 7) but not for positive psychotic experiences in the UK Biobank. To aid the comparison between the three psychotic experiences cohorts, we performed SNP annotations and gene mapping analyses on all psychotic experiences summary statistics using the same quality control procedures, methods and parameters within the FUMA pipeline (10) as follows: LD independent lead SNPs were identified at $p < 1 \times 10^{-5}$ for PE, at $p < 1 \times 10^{-6}$ for MDD, and at $p < 1 \times 10^{-8}$ for schizophrenia and bipolar disorder (p-value thresholds were set to allow for more than 20 independent SNPs to be analysed) within a 250kb window at $r^2 < 0.1$ based on LD structure in the 1000 Genomes phase 3 reference panel for individuals of European decent.

Annotation of functional consequences associated with independent lead SNPs and SNPs obtained from the reference panel that are in LD with independent SNPs (at $r^2 \geq 0.6$) was performed using ANNOVAR (11) (based on Ensembl genes build version 92) whilst excluding the extended MHC region (25,000,000-35,000,000). ANNOVAR is a software tool used to identify whether SNPs are associated with protein coding or amino acid changes. Annotation are based on several sources of information such as gene or splicing site locations, mRNA sites, genomic region-based information such as conserved regions and predicted transcription factor binding sites, stable RNA secondary structures or microRNA target sites. ANNOVAR offers the utility to use several public databases for a range of functional annotations as well as options on which to filter variants, such as SIFT scores for non-synonymous mutations. Based on user-defined gene definition databases like Ensembl, ANNOVAR annotates each variant to indicate its position in relation to genes (for instance, whether the variant is exonic, intronic, within a splicing site, upstream or downstream from a gene). For non-synonymous single nucleotide variants or indels, amino acid changes are also annotated. Precomputed functional importance scores, such as CADD scores (12), that indicate how likely a variant would have deleterious consequences, can also be annotated to variants. Based on these variant annotations, ANNOVAR offers the option to automate the process of gene mapping according to user-defined parameters.

Mapping of variants to the most likely causal genes was performed by employing a combination of positional mapping, expression quantitative trait loci (eQTL) mapping and 3D chromatin interaction mapping using the following parameters. Gene mapping was performed on lead independent SNPs and SNPs from the 1000 Genomes reference panel for individuals of European descent that were in LD with lead SNPs at $r^2 > 0.6$. For positional mapping, variants located within 10kb of known gene regions were mapped to genes if likely to be deleterious based on a CADD score ≥ 12.37 (12). eQTL mapping of SNPs to genes were performed based on significant eQTL associations at a false discovery rate (FDR) < 0.05 obtained from 13 brain regions from GTEx v7 brain tissue repository and 10 from GTEx v6 (13, 14). SNPs were mapped to genes based on significant chromatin interactions obtained from high-resolution HiC datasets for fetal and adult human brain samples (15) and for the dorsolateral prefrontal cortex and the hippocampus from GSE87112 at the recommended FDR of $p < 1 \times 10^{-6}$ 250kb upstream and 500kb downstream from the transcription start site (16). Promoter and enhancer regions were annotated from the Roadmap 111 epigenomes brain tissue for 13 brain regions (17, 18). Additionally, parameters in FUMA was set to map variants within protein-coding regions only.
